# Gluconeogenesis plays a critical role in mycobacterial biofilm formation

**DOI:** 10.1101/2025.03.01.640511

**Authors:** Shweta Singh, Sapna Bajeli, Ashwani Kumar

## Abstract

Gluconeogenesis is an important pathway for bacteria and humans as it maintains the glucose levels in the system. These glucose molecules can be utilized as substrates for several cellular functions. Gluconeogenesis genes are important for the intracellular survival and pathogenicity of *Mycobacterium tuberculosis* (Mtb). A few reports have shown the role of gluconeogenesis in biofilm formation in other bacteria. Similarly, we utilized gluconeogenesis genes and explored their role in biofilm formation in Mtb. For this study, we have taken the pyruvate carboxylase (*pca*) gene, which catalyzes the conversion of pyruvate into oxaloacetate and diverts the carbon flux into the gluconeogenesis pathway. We observed that the transposon mutant of *pca* was deficient in pellicle and submerged biofilm formation and was defective in colony morphology. This phenomenon was recovered upon either gene complementation or external addition of glucose into the medium. In addition to glucose, we also observed the regaining of biofilm phenomenon in the case of pyruvate supplementation. These observations led us to hypothesize that the glucose molecules generated from the gluconeogenesis pathway can be utilized to generate carbohydrates or polysaccharides, such as cellulose, which can be integrated into the extracellular polymeric substances of the biofilm matrix. Cellulose, whose primary subunit is glucose, can be synthesized from these generated glucose molecules, thereby integrating into the biofilm matrix of mycobacteria. This study provides a novel mechanism of cellulose production through a non-canonical pathway in Mtb, which is important in the biofilm formation of this pathogenic bacterium.

## Introduction

*Mycobacterium tuberculosis* (Mtb) is the causative agent of the highly infectious disease tuberculosis. Infection is transmitted by aerosols carrying a low bacterial number. These aerosols can be inhaled by human beings. Mtb, in many cases, is recognized by our immune system, monitoring the replication of pathogens^1^. Bacterial metabolism has been studied in-depth in the gram-positive intracellular pathogen, i.e., Mtb, as it is linked to its disease progression. As Mtb primarily resides within the phagosome of infected macrophages, it travels to multicellular caseous granuloma in the lung. The bacteria lay in a quiescent state, awaiting activation and disease progression. Mtb relies on host lipids, fatty acids, carbon sources, and cholesterol for energy sources to survive under these conditions^2^. Mtb is dependent on the gluconeogenesis pathway for its metabolic and virulence requirements. The statement is supported by the fact that pyruvate carboxykinase (*pckA*) is required for its full virulence^3^. In addition to *pckA*, genes *glpX,* and *gpm2*, which encode for fructose bisphosphatase in the Mtb gluconeogenetic pathway, are also required for virulence. Mutation of the above-mentioned genes leads to disruption of gluconeogenesis, resulting in attenuation of Mtb in *in vivo* models^4^. Gluconeogenesis is the pathway by which glucose is generated from non-carbohydrate carbon sources such as pyruvate, glycerol, acetate, lactate, and other glucogenic amino acids **(Fig. 1)**. This energy-intensive procedure generates carbohydrates for the bacteria^5^. It is also known as the reverse of the glycolysis pathway, as the enzymes in glycolytic and gluconeogenic pathways are the same, catalyzing readily reversible reactions^6^. The only irreversible reaction in the gluconeogenesis pathway is catalyzed by fructose-1, 6-bisphosphatase. Phosphoenolpyruvate is usually the entry point into the gluconeogenesis pathway, but it can vary depending on the carbon substrate^7^.

**Fig. 1:**
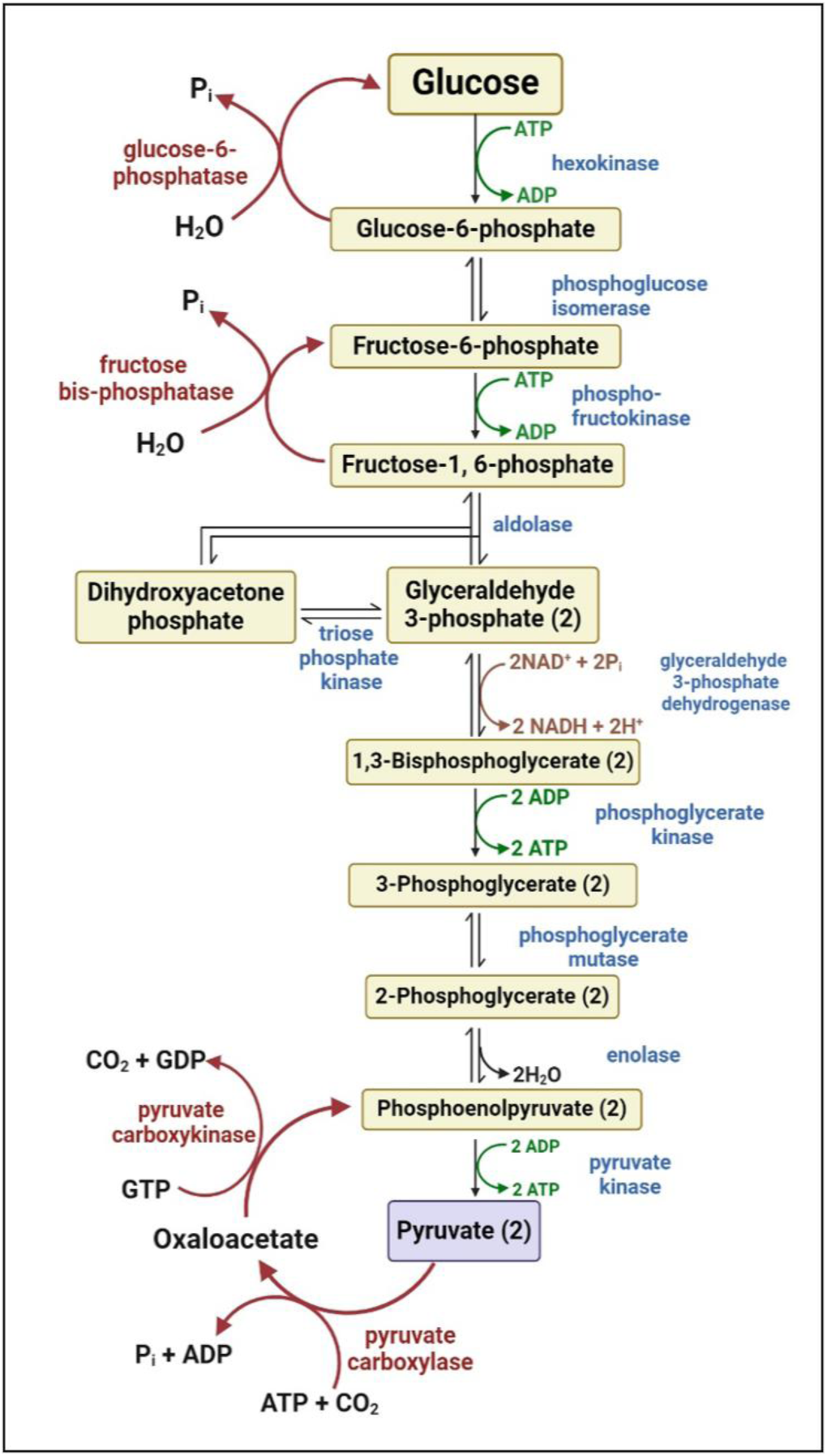
Schematic representation of gluconeogenesis/glycolysis pathway. The gluconeogenesis pathway is typically the reverse of glycolysis, with few key enzymatic steps involved in producing glucose from non-carbohydrate carbon sources, such as pyruvate or phosphoenolpyruvate. The enzymes in red are involved in the gluconeogenesis pathway, whereas those in blue are involved in the glycolysis pathway (Created in BioRender. Kumar, A. (2025) https://BioRender.com/c58q988).

Recent research on *Saccharomyces cerevisiae*, *Staphylococcus aureus*, *Salmonella enterica* serovar Typhimurium, *Vibrio cholerae*, and others have demonstrated that gluconeogenesis intermediates are necessary for extracellular polymeric substances (EPS) synthesis and biofilm development^8^. In the case of *S. cerevisiae*, a comparative RNA seq analysis was performed between its biofilm vs planktonic state, which showed a shift in the gluconeogenesis pathway during the early attachment phase of biofilm formation, whereas a shift towards glycolytic pathway during the maturation of biofilms. This observation showed that gluconeogenesis might have a role to play in biofilm formation^8^. In *S. aureus*, the bacterium produces itaconate, which blocks the glycolysis pathway and exhibits increased carbon flux through gluconeogenesis, to produce biofilm. An increased expression of pyruvate carboxylase (*pca*) was observed, a precursor of gluconeogenesis (an enzyme involved in oxaloacetate generation).

Itaconate treatment diverted the carbon flux into EPS production and enhanced the biofilm formation of *S. aureus*^9^. Another study in *Salmonella enterica* serovar Typhimurium showed that c-di-GMP can rewire the carbon flux into the gluconeogenesis pathway, which in turn provides the substrates for polysaccharide synthesis required in biofilm formation. At high concentrations (concs.) of c-di-GMP, an increased expression of *fbp*, *ppsA*, and *pck* genes, encoding for critical enzymes in the gluconeogenesis pathway, was observed. *Salmonella* harnesses the c-di-GMP molecule and acts as a double-edged sword by attenuating bacterial motility to promote biofilm formation and also coordinates carbon metabolism to gluconeogenesis to provide substrates for biofilm formation^10^. *Candida albicans* showed that the bacterium switches between the glycolysis/gluconeogenesis pathway during mature biofilm formation to provide hexose by-products that are utilized to facilitate biofilm formation and energy requirements^11^. Gluconeogenesis is also important for motility and biofilm formation in *Vibrio cholerae*. As the gluconeogenic substrates are important for the growth of *V. cholerae* in *in vivo* conditions^12^.

As described in the previous section, gluconeogenic substrates are a vital requirement for several pathogenic and non-pathogenic bacterial biofilm formation^8–10,12^. These bacteria harness the gluconeogenesis pathway to generate EPS components and polysaccharide biosynthesis to integrate into the biofilm matrix. Previously known literature highlights the importance of gluconeogenesis in Mtb metabolism and virulence. Although, the involvement of gluconeogenesis in mycobacterial biofilm formation remains unexplored. In this study, we have checked the role of gluconeogenesis genes in mycobacterial biofilm formation. Ever since the presence of cellulose was demonstrated in mycobacterial biofilms, the presence of biosynthetic genes required for cellulose production has been explored^13–15^. Since glucose is the primary subunit required for cellulose production. We hypothesize the glucose required for cellulose production could be generated through increased carbon flux into the gluconeogenesis pathway. Like the other bacteria, Mtb might also increase gluconeogenesis pathway genes for EPS and polysaccharide synthesis during biofilm formation.

## Results

### Impact of different carbon sources in biofilm formation of Mtb H37Rv

The role of carbon sources in biofilm formation has been previously checked in pathogenic microorganisms such as *P. aeruginosa*, *M. avium*, *C. albicans*, and *C. glabrata,* etc^16–19^. These reports have demonstrated that the addition of carbon sources, such as glucose, acetate, lactate, arabinose, etc., can influence the biofilm formation of the microorganism. It has been observed that glucose can generally enhance biofilm formation, as it can contribute to the carbohydrate content of EPS in biofilms.

Using a similar hypothesis, we have used different carbon sources previously used^20^ to determine the effect on Mtb’s growth. For this study, we have taken carbon sources such as glucose, pyruvate, acetate, succinate, malate, and oxaloacetate. Typically, Sauton’s media contains glycerol as the primary carbon source for utilization during biofilm formation. Here, we replaced glycerol with the above-mentioned carbon sources, i.e., glucose conc. ranging (10 mM-200 mM), pyruvate conc. ranging (2 mM-50 mM), acetate conc. ranging (5 mM- 100 mM), succinate conc. ranging (0.5 mM- 5 mM), malate conc. ranging (0.5 mM- 6 mM), and oxaloacetate conc. ranging (1 mM- 8 mM).

In the case of succinic and malic acid, the biofilm capability was hampered, and the bacteria could tolerate 5 mM and 6 mM of their conc., respectively **(Fig. 2a, b)**. In addition, we observed that in the case of oxaloacetic acid, the strain could tolerate up to 8 mM of its conc. and was able to form biofilm in its presence, although these biofilms were very thin, unlike the control (glycerol), which led to the formation of mature pellicle biofilms **(Fig. 2c)**. These observations were supported by quantitative crystal violet (CV) assays, which showed that glycerol panel showed higher biofilm formation as compared to other carbon sources, and there was a statistically significant difference between the control and succinic, malic, and oxaloacetic acid panels **(Fig. 2d-f)**. Acetic acid and pyruvate were able to support the biofilm formation, but the biofilm was very thin and fragile in comparison to the control panel and tolerated up to 150 mM and 70 mM of their conc., respectively **(Fig. 2g, i)**. Importantly, glucose supports the formation of mature and fully developed biofilms, which are comparable to the biofilms formed in the presence of glycerol **(Fig. 2h)**. Upon biofilm quantitation using CV assay, a statistically significant difference was observed between the glycerol panel and different concs. of acetic acid and pyruvate **(Fig. 2j, l)**. In the case of glucose, the concentrations, i.e., 50 mM, 100 mM, and 150 mM, showed elevated biofilm formation, and the data was similar to that of the control group. 100 mM glucose could substantially support biofilm formation, and 150 mM glucose can help form mature and fully developed biofilm similar to glycerol **(Fig. 2k)**. These results suggested that glucose as a carbon source could facilitate biofilm formation apart from glycerol.

**Fig. 2:**
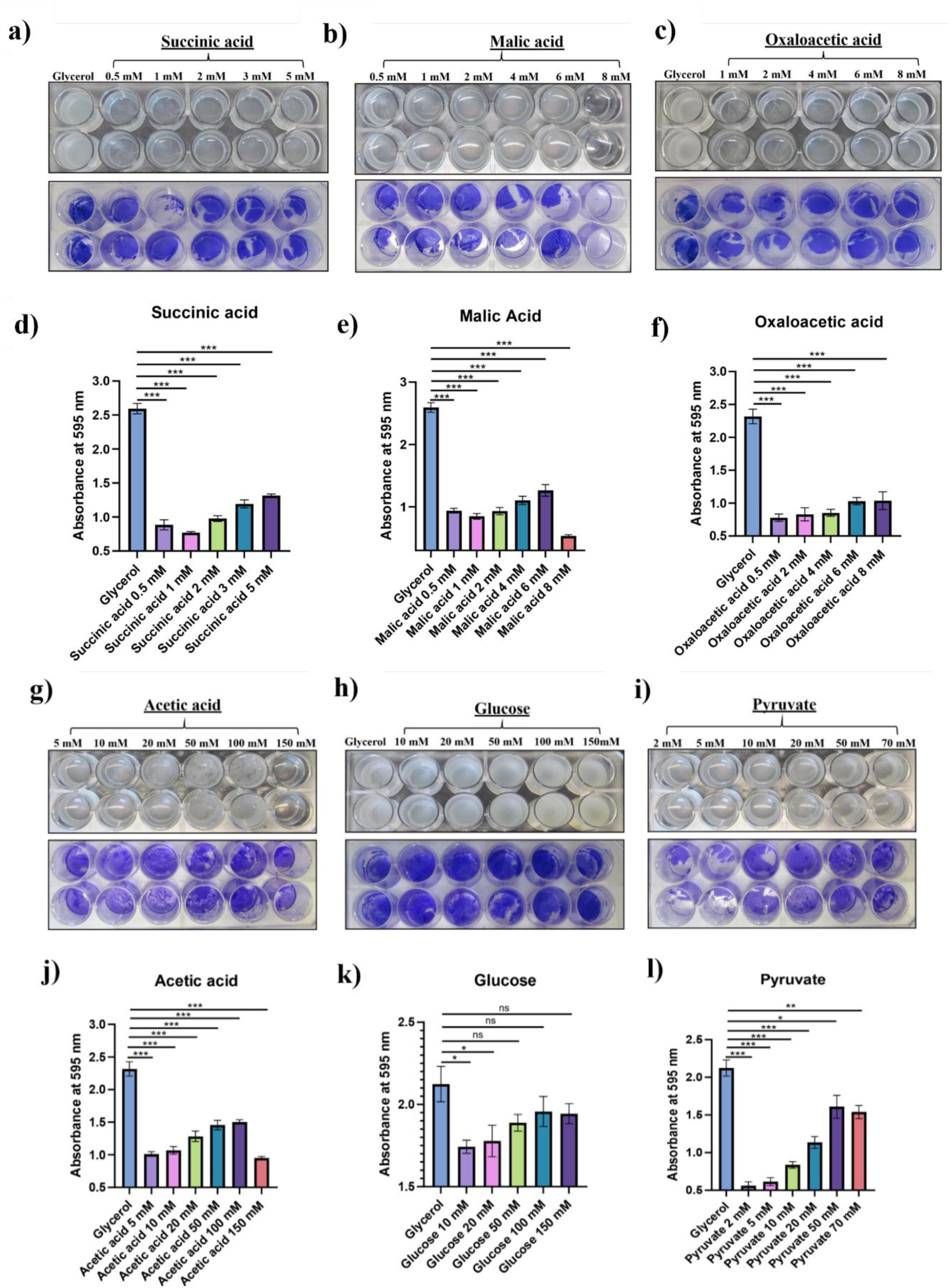
In the absence of glycerol, glucose, and pyruvate can restore the biofilm formation in Mtb H37Rv. Representative image of Mtb H37Rv pellicle biofilm formation in 24-well plates with glycerol in the first well taken as a positive control, with other wells containing **a)** differing concs. of succinic acid ranging from 0.5 mM to 5 mM. **b)** differing malic acid concentrations ranging from 0.5 mM to 8 mM, **c)** differing concs. of oxaloacetic acid ranging from 0.5 mM to 8 mM. Graph showing CV assay of pellicle biofilm formed in 24-well plates with **d)** glycerol and succinic acid carbon sources, **e)** glycerol and malic acid as carbon sources, **f)** glycerol and oxaloacetic acid as carbon sources. Representative image of Mtb H37Rv pellicle biofilm formation in 24-well plates with **g)** differing concs. of acetic acid ranging from 5 mM to 150 mM **h)** differing concentrations of glucose ranging from 10 mM to 150 mM **i)** differing concs. of pyruvate ranging from 2 mM to 70 mM. Graph showing CV assay of pellicle biofilm formed in 24-well plates with **j)** glycerol and acetic acid as carbon sources, **k)** glycerol and glucose as carbon sources, **l)** glycerol and pyruvate as carbon sources. The experiments shown in **a-c** and **g-i** are performed in duplicates. Data represent mean±SEM from three independent experiments. GraphPad Prism 8 was used to plot the column bar graphs. Statistical significance was determined using Student’s t-test, * indicates a p-value of <0.05, ** indicates a p-value of <0.005, *** indicates a p-value of <0.0005.

### *pca::Tn* is defective for pellicle biofilm formation

After establishing that glucose is important for biofilm formation, we next aimed to explore the genes involved in glucose formation, i.e., gluconeogenesis. To check whether the gluconeogenesis node has any role in biofilm formation, we have utilized a transposon (Tn) mutant of the *pca* gene, a probable pyruvate carboxylase, and checked its biofilm-forming capability. We have also taken Mtb CDC1551 as the wild-type strain, as the Tn mutant of *pca* was in this strain. When we subjected these strains to pellicle biofilm formation, we observed that *pca::Tn* could not form pellicle biofilm compared to the wild-type strain **(Fig. 3a)**, suggesting its plausible role in biofilm formation. Spot assays of these strains on 7H11 oleic acid-albumin-dextrose-sodium chloride (OADC) media and 7H11OADC containing Congo Red (CR) and Coomassie Brilliant Blue dyes (CBB) media showed that the Mtb CDC1551 strain was able to pick up the CR and CBB dyes, demonstrating the presence of carbohydrates and proteins in the colony. Meanwhile, in the case of *pca::Tn*, the colony was defective in the color uptake of the CR and CBB dyes, showing the lack of carbohydrates and protein in the colony phenotype **(Fig. 3b)**. We also evaluated the submerged biofilm formation capability of *pca::Tn* and Mtb CDC1551 strain in the presence of 6mM dithiothreitol (DTT)^15^. It was observed that the *pca::Tn* strain was defective in submerged biofilm as compared to the Mtb CDC1551 strain **(Fig. 3c)**. Overall, these results suggest that the absence of the *pca* gene leads to defects in biofilm formation and color uptake by the macrocolony, indicating its importance in the biofilm formation mechanism in mycobacteria. In addition to *pca*, we have also utilized *pckA* mutant and checked for its biofilm-forming capability and ability to uptake color by the macrocolony. The strain was able to form biofilm but appeared to be less corded than the wild-type strain **(Fig. 3a)**. Similarly, spot assay results showed that in 7H11OADC media, the colony was corded, although less than Mtb CDC1551, and was able to pick the color on 7H11OADC CR-CBB media **(Fig. 3b)**. The *pckA* mutant was quite similar to the Mtb CDC1551 strain. Since the *pckA* mutant showed a similar phenotype to Mtb CDC1551, we have proceeded with the *pca::Tn* strain in further experiments.

**Fig. 3:**
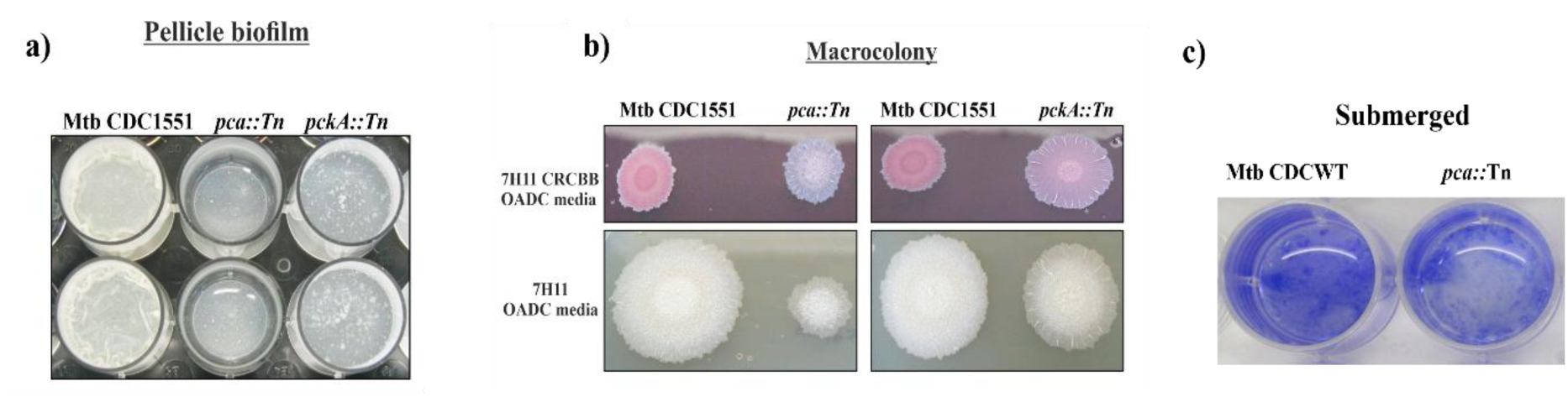
***pca::Tn* shows a defect in the pellicle and submerged biofilm and colony morphology. a)** Differences in pellicle biofilm formation in 24-well plates between Mtb CDC1551, *pca::Tn*, and *pckA::Tn* strains, where *pca:: Tn* shows defective biofilm formation. *pckA::Tn* showed pellicle biofilm formation with fewer cordings as compared to the Mtb CDC1551 strain. **b**) Differences in colony morphology between Mtb CDC1551, *pca::Tn*, and *pckA::Tn* strains on 7H11OADC and 7H11OADC CR-CBB agar plates. *pca::Tn* colony is defective in color uptake of Congo red (CR) and Coomassie brilliant blue (CBB) dyes, depicting the absence of polysaccharide and curli protein content in the colony. It also shows smaller colonies with fewer cordings than the Mtb CDC1551 strain. *pckA::Tn* shows slight differences in colony morphology and grew similarly to the Mtb CDC1551 strain. **c)** Differences in the submerged biofilm formation between the *pca::Tn* and Mtb CDC1551 strain upon 6 mM DTT induction.

### Impact of different carbon sources in biofilm formation of Mtb CDC1551 and *pca::Tn* strain

As described previously, we also checked the effect of different carbon sources on Mtb CDC1551 and *pca::Tn* strain. Glycerol was replaced with carbon sources, i.e., succinate, malate, oxaloacetate, acetate, glucose, and pyruvate.

Determination of the pellicle biofilm formation was observed visually as well as quantitated through CV assay. In the Mtb CDC1551 strain, glycerol containing Sauton’s media was taken as a positive control for the experiments. It was observed that glycerol could support the formation of mature pellicle biofilms, succinic and malic acid were taken as the sole carbon sources, the capability to form biofilm was hampered, and the bacteria could tolerate 5 mM and 6 mM of their conc., respectively **(Fig. 4a, b)**. Whereas, in the case of oxaloacetic acid the strain could tolerate up to 6 mM of its conc. and cells were found to be settled at the bottom **(Fig. 4c)**. A similar experimental setup was used for the *pca::Tn* strain, and its biofilm formation ability was checked under different carbon sources. Here, in the glycerol panel (control), the *pca::Tn* strain is unable to form a biofilm, and upon addition of different carbon sources such as succinic acid, malic acid, and oxaloacetic acid failed to restore the wild-type phenomenon in the mutant strain as depicted visually **(Fig. 4d-f)**. This phenomenon was also quantitated through CV assay for both the strains, and comparative graphs were plotted for three different carbon sources, i.e., succinic acid, malic acid, and oxaloacetic acid **(Fig. 4g-i)**. It was evident from the graphs that in the case of glycerol, *pca::Tn* was defective in biofilm formation compared to Mtb CDC1551. These three carbon sources could not facilitate the biofilm formation for both Mtb CDC1551 and *pca::Tn* strains.

**Fig. 4:**
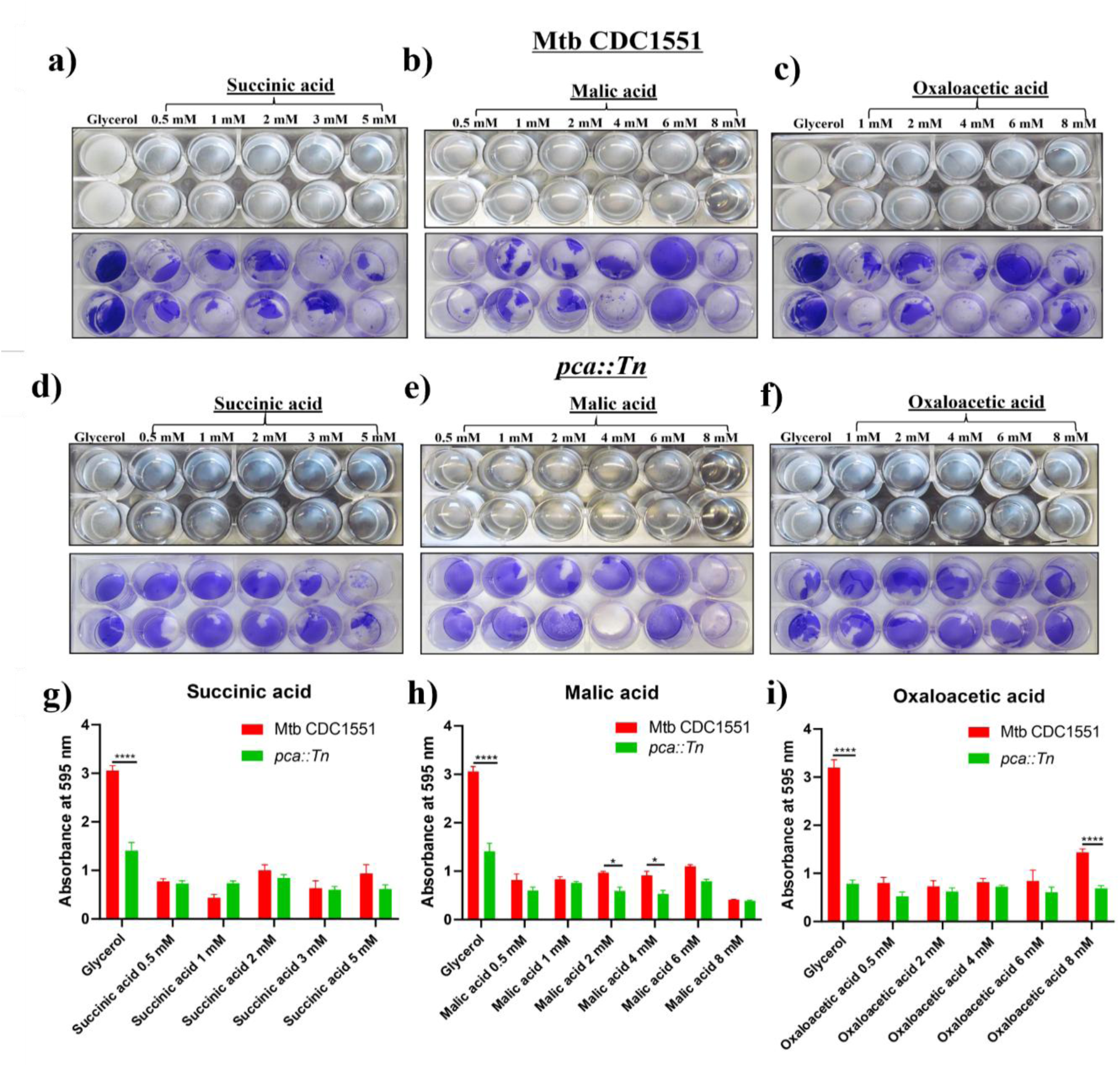
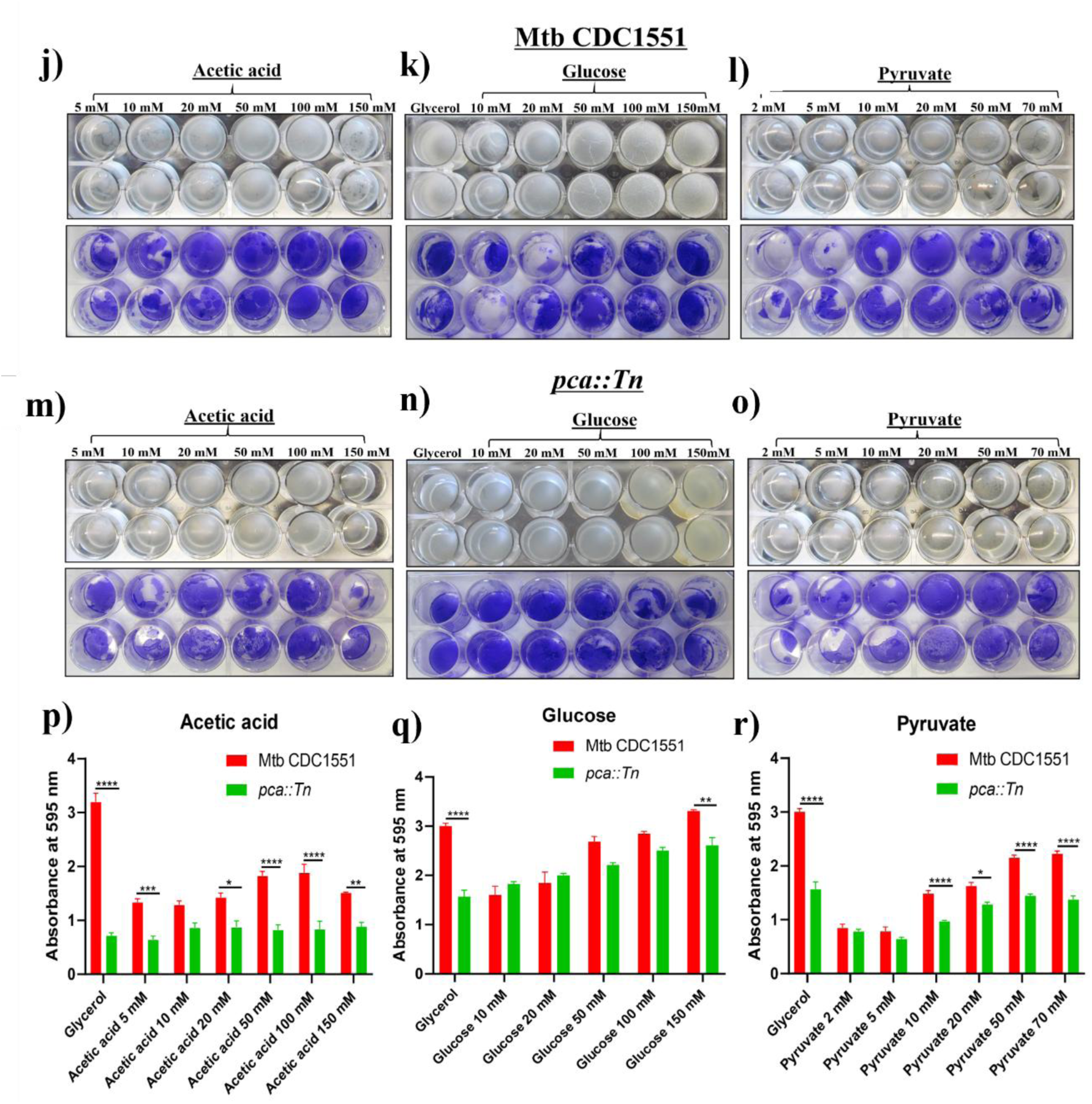
In the absence of glycerol, glucose, and pyruvate can restore the biofilm formation in Mtb CDC1551 and *pca::Tn* strains. Representative image of Mtb CDC1551 pellicle biofilm formation in 24-well plates with glycerol taken as a positive control, with other wells containing **a)** first well glycerol, rest differing concs. of succinic acid ranging from 0.5 mM to 5 mM **b)** differing concentrations of malic acid ranging from 0.5 mM to 8 mM **c)** first well glycerol, rest differing concs. of oxaloacetic acid ranging from 0.5 mM to 8 mM **j)** differing concs. of acetic acid ranging from 5 mM to 150 mM **k)** first well glycerol, rest differing concs. of glucose ranging from 10 mM to 150 mM **l)** differing concs. of pyruvate ranging from 2 mM to 70 mM. Representative image of *pca::Tn* pellicle biofilm formation in 24-well plates with glycerol as a positive control, with the rest of the wells containing **d)** first well glycerol, differing concs. of succinic acid ranging from 0.5 mM to 5 mM **e)** differing concentrations of malic acid ranging from 0.5 mM to 8 mM **f)** first well glycerol, differing concs. of oxaloacetic acid ranging from 0.5 mM to 8 mM **m)** differing concs. of acetic acid ranging from 5 mM to 150 mM **n)** first well glycerol, rest differing concentrations of glucose ranging from 10 mM to 150 mM **o)** differing concs. of pyruvate ranging from 2 mM to 70 mM. Graph showing CV assay of pellicle biofilm formed in 24-well plates with different carbon sources for both Mtb CDC1551 and *pca::Tn* strains. **g)** glycerol and succinic acid, **h)** glycerol and malic acid, **i)** glycerol and oxaloacetic acid, **p)** glycerol and acetic acid, **q)** with glycerol and glucose, **r)** glycerol and pyruvate. The experiments shown in **a-f** and **j-o** are performed in duplicates. Data represent mean±SEM from three independent experiments. GraphPad Prism 8 was used to plot the column bar graphs. Statistical significance was determined using Two-way ANOVA, * indicates a p-value of <0.05, ** indicates a p-value of <0.005, *** indicates a p-value of <0.0005, **** indicates a p-value of <0.0001.

In the case of acetic acid and pyruvate, the Mtb CDC1551 strain was able to form the biofilms, but the biofilms were fragile in comparison to the glycerol panel and tolerated up to 150 mM and 70 mM of their conc., respectively. Whereas, in the case of glucose, the bacteria showed the best complementation with respect to the control, i.e., glycerol **(Fig. 4j-l)**. The pellicle biofilm structure was sturdy and possessed cordings similar to the glycerol panel. In the case of the *pca::Tn* mutant strain, acetic acid failed to restore the wild-type phenomenon, whereas glucose enabled the bacilli to form biofilms similar to the Mtb CDC1551 in the glycerol panel **(Fig. 4m)**. Presence of glucose conc. 50 mM, 100 mM, and 150 mM in the culture medium enhanced biofilm formation and the presence of cordings in the biofilm depicting its sturdiness **(Fig. 4n)**. However, to a lesser extent, pyruvate also enabled the formation of biofilm phenomenon in the *pca::Tn* mutant strain at concentrations 20 mM, 50 mM, and 70 mM, as depicted visually **(Fig. 4o)**. This phenomenon was quantitated through CV assay for both strains, and comparative graphs were plotted for three different carbon sources, i.e., acetic acid, glucose, and pyruvate **(Fig. 4p-r)**. It was evident from the graphs that in the case of glycerol, *pca::Tn* was defective in biofilm formation compared to Mtb CDC1551. Only glucose could provide the best complementation of the biofilm defect in the case of Mtb CDC1551 and *pca::Tn* strains, as depicted through CV assay **(Fig. 4p, r)**. A comparable biofilm formation was observed for both strains, with Mtb CDC1551 showing even better biofilms at 150 mM glucose conc. Acetic acid and pyruvate also facilitated biofilm formation but to a lesser extent. These results suggested that, apart from glycerol, glucose enabled the biofilm phenotype in the absence of any other carbon source, suggesting that glucose and glycerol are important carbon sources required for biofilm formation.

### *pca::Tn* strain is defective in growth and is rescued upon either complementation or addition of glucose

The above-described experiments suggest that the *pca* gene is important for mycobacterial biofilm formation and colony morphology. To ensure that the *pca* gene is indeed responsible for the biofilm phenomenon, we complemented the mutant strain by reintroducing the *pca* gene into the genome under the control of the constitutive hsp60 promoter. The vector control strain was also constructed where only a vector with no gene was inserted within the *pca::Tn* strain. To investigate the complementation of the *pca* gene in the *pca::*comp strain, we checked mRNA levels in the Mtb CDC1551, *pca::Tn*, *pca::Tn* VC, and *pca::comp* strains. It was observed that the complemented strain showed elevated mRNA expression similar to the wild-type strain. Meanwhile, the mutant and vector control strains showed lower mRNA levels. The difference in expression levels of *pca::Tn* between WT was significant. In contrast, the fold difference in mRNA level between WT and the complemented strain remains nonsignificant (not indicated) **(Fig. 5a)**. These four strains, Mtb CDC WT, *pca::Tn* (mutant), *pca::Tn VC* (vector control strain), and *pca::Tn comp* (complemented strain), were checked for their *in vitro* growth pattern in 7H9 glycerol media. It was observed that the mutant strain was found to be defective in growth as compared to the wild-type strain. Whereas the complemented strain was found to regain the defect and grew similar to the wild-type strain. *pca::Tn VC* was taken as a vector control strain that grew similarly to the mutant strain *pca::Tn* **(Fig. 5b)**. A significant difference was observed between the WT and *pca::Tn* strains at several time points. Thereafter, we grew the strains in a 7H9 medium containing glucose as a carbon source and checked for the *in vitro* growth pattern. We observed that all four strains grew similarly, and the growth defect in the mutant strain was restored upon the glucose supplementation **(Fig. 5c, d)**. There was no difference between the growth rate of strains, and they were growing similarly to the wild type in a medium containing 100 mM and 150 mM glucose concentrations as carbon sources.

**Fig. 5:**
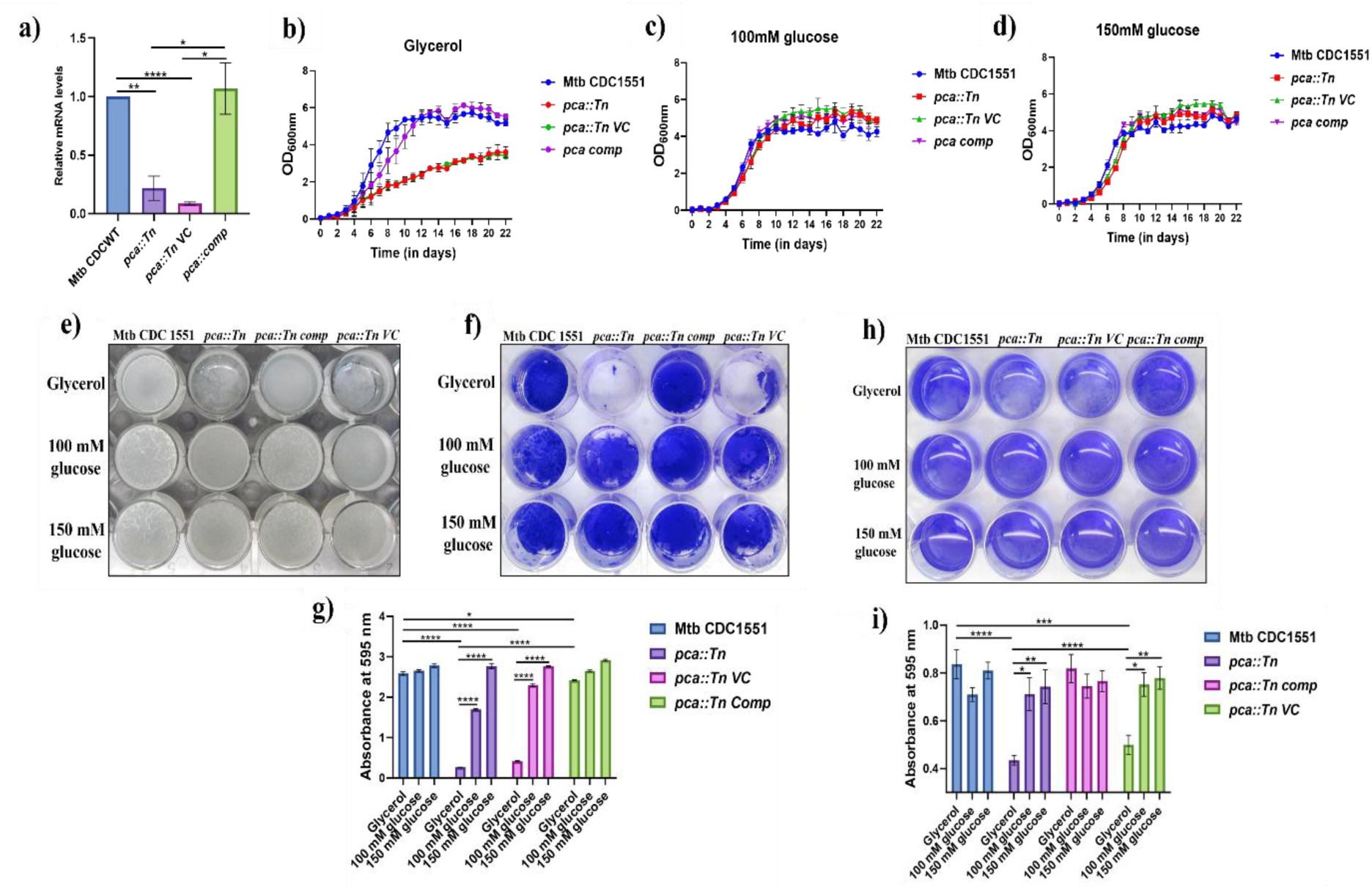
The addition of glucose or *pca* gene complementation restores the growth defect, pellicle, and submerged biofilm phenotype in the mutant strain. a) Column bar graph showing the relative mRNA levels between the four strains. The complemented strain showed elevated mRNA expression similar to the wild-type strain. Meanwhile, the mutant and vector control strains showed lower mRNA levels. Growth profile of Mtb CDC1551, *pca::Tn*, *pca::Tn VC*, and *pca::Tn comp* where **b)** glycerol, **c)** 100 mM glucose, and **d)** 150 mM glucose were utilized as a carbon source. **e)** Representative image of pellicle biofilm formation in 24-well plates of Mtb CDC1551, *pca::Tn*, *pca::Tn VC*, and *pca::Tn comp* strains with glycerol and glucose as carbon sources. **f)** Representative image of CV assay performed on pellicle biofilm setup. **g)** Graph showing CV assay of pellicle biofilm of Mtb CDC1551, *pca::Tn*, *pca::Tn VC*, and *pca::Tn comp* strains with glycerol and glucose as carbon sources. **h)** Representative image of submerged biofilm formation in 24-well plates of Mtb CDC1551, *pca::Tn*, *pca::Tn VC*, and *pca::Tn comp* strains; 6 mM DTT for 29 hr was given to induce biofilm formation. Glycerol and glucose were utilized as carbon sources. **i)** Graph showing CV assay of submerged biofilm of Mtb CDC1551, *pca::Tn*, *pca::Tn VC*, and *pca::Tn comp* strains with as carbon source. Data represent mean±SEM from three biological triplicates, each with two technical repeats **(a)**. The XY graphs were plotted using GraphPad Prism 8. Data represent mean±SEM from three independent experiments **(b-d)**. Statistical significance was determined using Unpaired t-test, * indicates a p-value of <0.05, ** indicates a p-value of <0.005, *** indicates a p-value of <0.001, **** indicates a p-value of <0.0001 **(a)**. Data represent mean±SEM from three independent experiments **(e, f, h)**. Statistical significance was determined using Two-way ANOVA, * indicates a p-value of <0.05, ** indicates a p-value of <0.005, *** indicates a p-value of <0.0005, **** indicates a p-value of <0.0001 **(g, i)**. GraphPad Prism 8 was used to plot the column bar graphs.

### Complementation of the *pca* gene or addition of glucose leads to restoration of the defect in pellicle biofilm phenotype in the *pca::Tn* strain

Next, we sought to check the biofilm-forming capacity of Mtb CDC1551, *pca::Tn*, *pca::Tn VC*, and *pca::Tn comp* strains upon gene complementation or in the presence of glucose. As predicted, we observed that the complemented strain was able to regain the biofilm formation ability in Sauton’s media. The complemented strain was able to show cording similar to the wild-type strain and was also able to grow normally in contrast to the mutant strain, which showed retarded growth on media plates. *pca::Tn VC* was taken as a control strain harboring only the plasmid without the gene behaving similarly to the *pca::Tn* mutant strain **(Fig. 5e, f)**. We sought to check whether the defect in biofilm formation could be restored upon adding glucose in the Sauton’s media, as described in the previous results. Toward this, we have taken two glucose concentrations, i.e., 100 mM and 150 mM, at which we observed the best phenotypic complementation. We observed that the addition of glucose concs. i.e., 100 mM and 150 mM conc. restore the biofilm defect in the *pca::Tn* strain and enhance biofilm formation in the wild type and the complemented strain, which otherwise can form biofilm in the control panel **(Fig. 5e, f)**. This phenomenon was quantitated using CV assay. The CV assay values were plotted, and the graph obtained showed a significant difference between the Mtb CDC1551 and *pca::Tn* mutant strain when grown upon glycerol as a carbon source. As the glucose was supplemented, the mutant strain showed biofilm formation comparable to the wild-type strain. The data represents the values obtained from three biological replicates **(Fig. 5g)**.

In addition to quantification via CV assay, we also imaged the pellicle biofilm formed by the above-mentioned strains via confocal laser scanning microscopy (CLSM). Pellicle biofilms were formed in 8-well chambered glass slides and stained the different mycobacterial strains using Auramine O dye and cellulose using IMT-CBD-mC stain, which specifically binds to cellulose binding domain (CBD)^14^. Here, we observed that the Mtb CDC1551 strain showed thicker and voluminous biofilms in contrast to the *pca::Tn* mutant and *pca::Tn VC* strain, where the biomass in the biofilm structure was lesser than in comparison with wild type and *pca::Tn comp* strain and also consisted of few gaps/spaces in between the biofilm structures **(Fig. 6a)**. Also, the cells in the biofilm were more scattered in the mutant and VC strain than the Mtb CDC1551, as visualized in the Z-stacks showing 3D structures with well-formed voluminous biofilm structures. Upon glucose supplementation in the medium at 100 mM and 150 mM glucose conc. it was observed that the mutant strain regained the phenomenon and formed thicker biofilms similar to the wild-type strain with no gaps/spaces in the structure. We also observed that IMT-CBD-mC staining was lesser in the mutant strain than in the control panel, whereas increased staining was visualized in the case of 100 mM and 150 mM glucose panels for all four strains **(Fig. 6b-c)**. Enhanced staining in the glucose panels was observed, indicating the enhanced cellulose/polysaccharide presence in the pellicle biofilms. As pellicle biofilms have been previously reported to have high polysaccharide content in their biofilms^13–15^. Representative images of the side view of the biofilm panels are shown in **Fig. 6d-f**, which showcases the biofilm thickness of different strains. Biomass and thickness analysis was performed for the panels described above, and the statistical difference was observed between Mtb CDC1551 and *pca::Tn*, *pca::Tn VC* strain, and between the *pca::Tn* glycerol and *pca::Tn* glucose panels, highlighting the role of glucose in biofilm formation **(Fig. 6g-h)**. Relative intensity was calculated between mcherry/Auramine O fluorescence to showcase the increasing mcherry intensity in panels with glucose concentrations **(Fig. 6i)**. The above-mentioned results suggested that glycerol/glucose is required for biofilm formation as these carbon sources provide energy as well as building blocks for EPS production in the biofilm. In the absence of *pca,* which is involved in the gluconeogenesis pathway, glucose production is hampered, resulting in deficient biofilm formation. Therefore, an external glucose supplementation is required to restore the phenomenon.

**Fig. 6:**
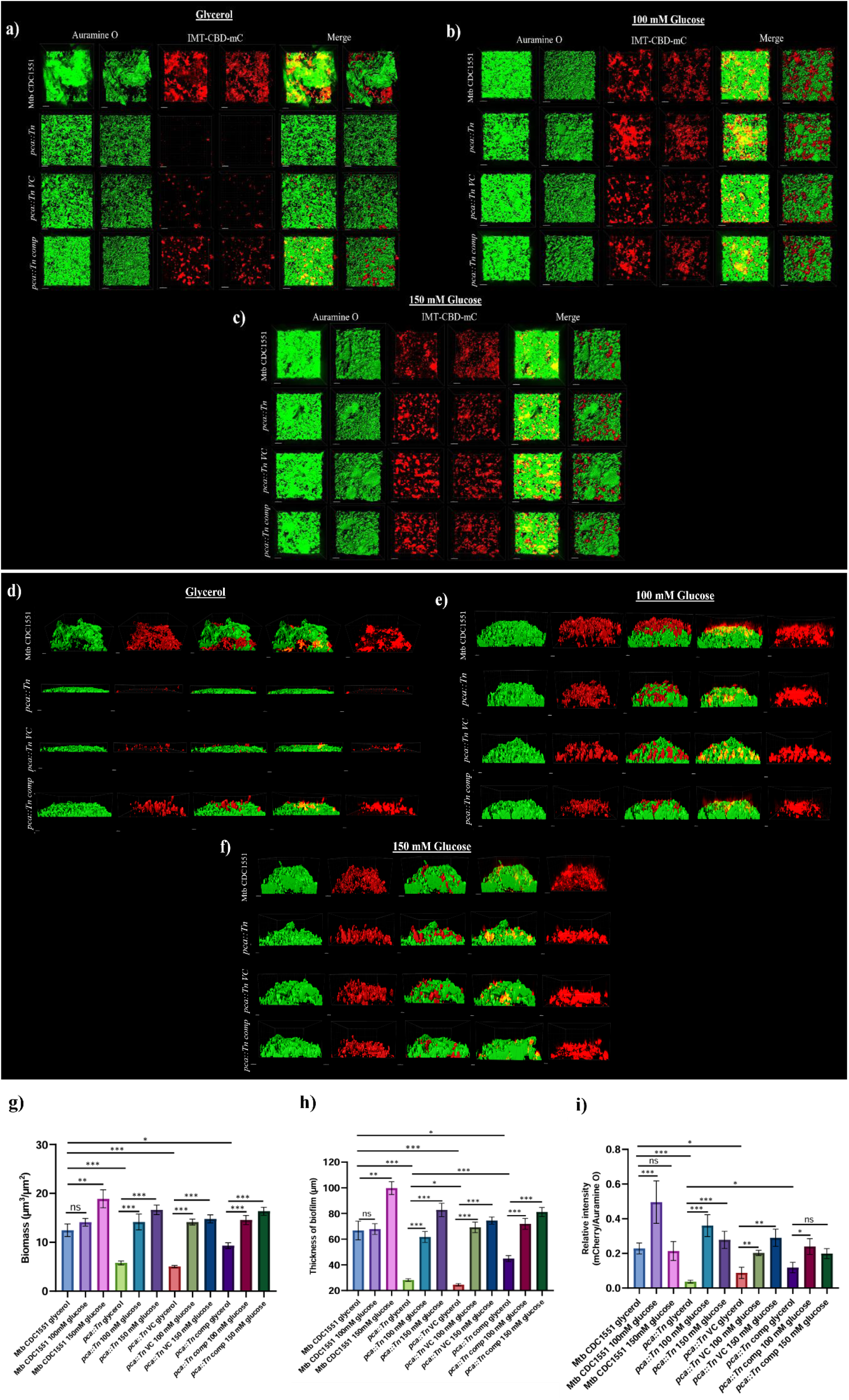
The addition of glucose regains the formation of thick, voluminous pellicle biofilm structures in the *pca::Tn* strain when visualized by CLSM. Representative 3D images of pellicle biofilm of all four strains i.e., Mtb CDC1551, *pca::Tn*, *pca::Tn VC*, and *pca::Tn comp* a) under glycerol-fed conditions. **b)** under 100 mM glucose-fed conditions. **c)** under 150 mM glucose-fed conditions. Representative side view images of pellicle biofilm of all four strains **d)** under glycerol-fed conditions. **e)** under 100 mM glucose-fed conditions. **f)** under 150 mM glucose-fed conditions. **g)** Biomass (in µm^3^/µm^2^), **h)** thickness (in µm), and **i)** relative intensity analysis of the biofilm structures. Biomass and thickness analysis was performed through the COMSTAT plugin in ImageJ software, and relative intensity was calculated through IMARIS version 8.9 software. Data represent mean±SEM from three independent experiments. GraphPad Prism 8 was used to plot the column bar graphs. Statistical significance was determined using Student’s t-test, * indicates a p-value of <0.05, ** indicates a p-value of <0.005, *** indicates a p-value of <0.0005, **** indicates a p-value of <0.0001.

### Complementation of the *pca* gene or addition of glucose leads to restoration of the defect in submerged biofilm phenotype in the *pca::Tn* strain

Our group has recently demonstrated that Mtb, Msmeg, and other mycobacterial species are capable of forming a biofilm upon exposure to thiol reductive stress (TRS)^14,15^. Thus, we analyzed the biofilm formation in *pca::Tn* strain upon TRS. To test this, logarithmic cultures of Mtb CDC1551, *pca::Tn*, *pca::Tn VC,* and its complemented strain were grown and then subjected to 6 mM DTT treatment. After exposure, DTT-induced biofilm formation was observed after 29 hours of exposure in Mtb CDC1551. Whereas the *pca::Tn* and *pca::Tn VC* lacked a rigid biofilm structure, the complemented strain was able to regain the phenomenon **(Fig. 5h)**. We sought to check whether the biofilm defect could be restored upon the addition of glucose in the 7H9+5% OADC medium. We observed that the addition of glucose concentrations, i.e., 100 mM and 150 mM, restores the biofilm formation capability in the *pca::Tn* strain and enhances biofilm formation in Mtb CDC1551 as well as the *pca::Tn comp* strain, which otherwise can form biofilm in the glycerol panel. This phenomenon was quantitated using CV assay **(Fig. 5h)**. The CV assay values were plotted, and the graph obtained showed a significant difference between the Mtb CDC1551 and *pca::Tn* and its vector strain when grown upon glycerol as a carbon source. The difference becomes non-significant upon the addition of glucose between the Mtb CDC1551 and *pca::Tn* strain. The data represent the values obtained from three biological replicates **(Fig. 5i)**.

In addition to quantification via CV assay, we also imaged the submerged biofilm formed by the above-mentioned strains via CLSM. Submerged biofilms were formed in 8-well chambered glass slides and stained the different mycobacterial strains using Auramine O dye, and cellulose using IMT-CBD-mC stain. Similar to the previous results **(Fig. 6)**, in the control panel (having glycerol as a carbon source), we observed that the Mtb CDC1551 wild-type strain showed thicker and fully matured biofilms in contrast to the *pca::Tn* mutant and *pca::Tn VC* strain where the biomass in the biofilm structure was lesser than in comparison with Mtb CDC 1551 and *pca::Tn comp* complemented strain and also consisted of few gaps/spaces in between the biofilm structures **(Fig. 7a)**. Whereas, in the case of glucose supplementation i.e., 100 mM and 150 mM glucose conc. it was observed that the mutant strain regained the capability to form the biofilm and was capable of forming thicker biofilms similar to the wild-type strain with no gaps/spaces in the structure. We also observed that IMT-CBD-mC staining was absent from the mutant strain in the case of the glycerol panel, whereas increased staining was visualized in the case of 100 mM and 150 mM glucose panels for all four strains **(Fig. 7b, c)**. Representative images of the side view of the biofilm panels are shown in **Fig. 7d-f**, which showcases the biofilm thickness of different strains. Biomass and thickness analysis were performed for the panels described above, and the statistical difference was observed between Mtb CDC1551 and *pca::Tn*, and *pca::Tn VC* and between the *pca::Tn* glycerol and *pca::Tn* glucose panels, highlighting the role of glucose in biofilm formation **(Fig. 7g, h)**. Relative intensity was calculated between mcherry/Auramine O fluorescence to showcase the increasing mcherry intensity in panels with glucose concentrations **(Fig. 7i)**. This suggested that in addition to pellicle biofilm, glycerol/glucose is also required to form submerged biofilms. These carbon sources provide energy and building blocks for EPS production in the biofilm.

**Fig. 7:**
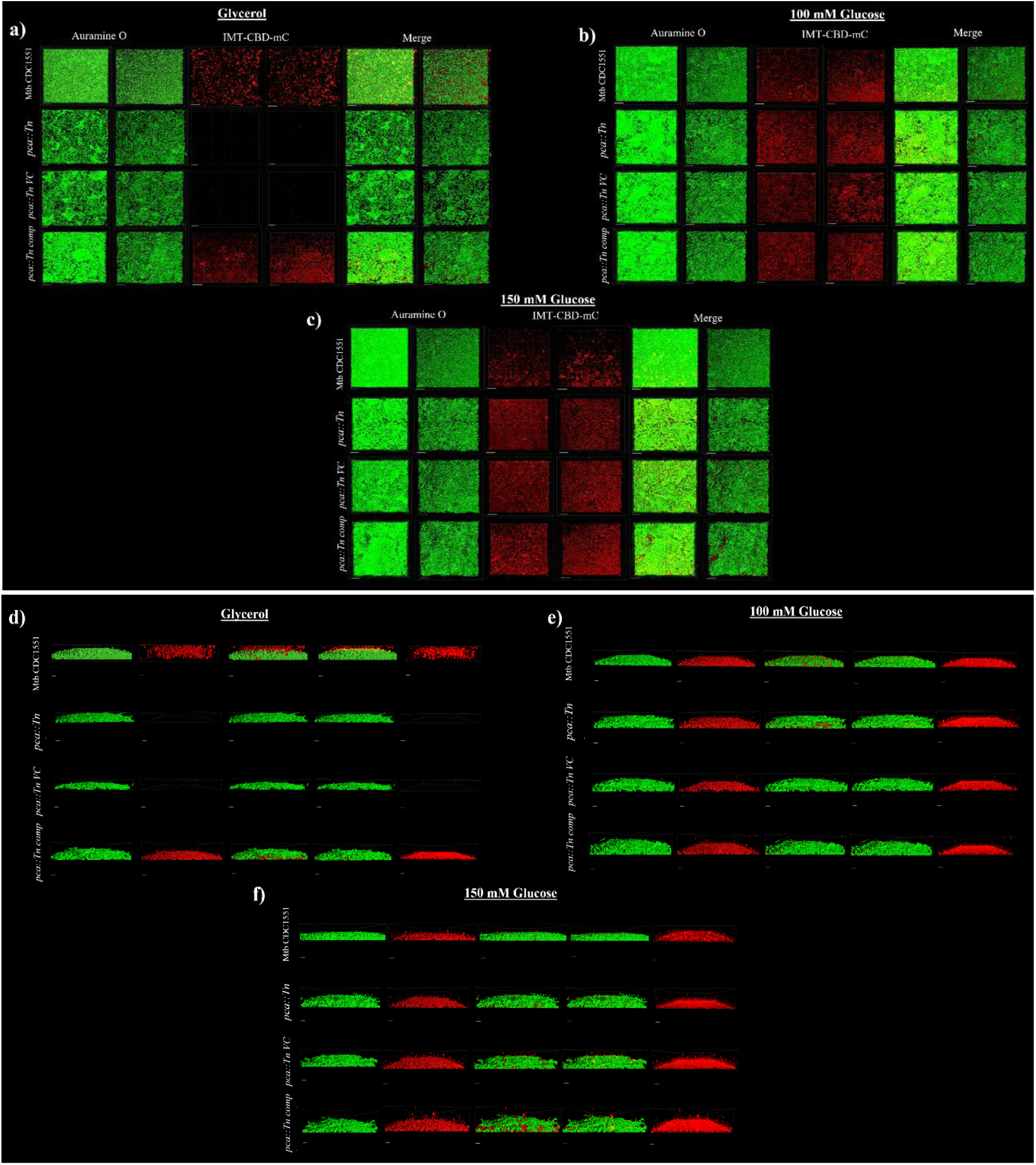
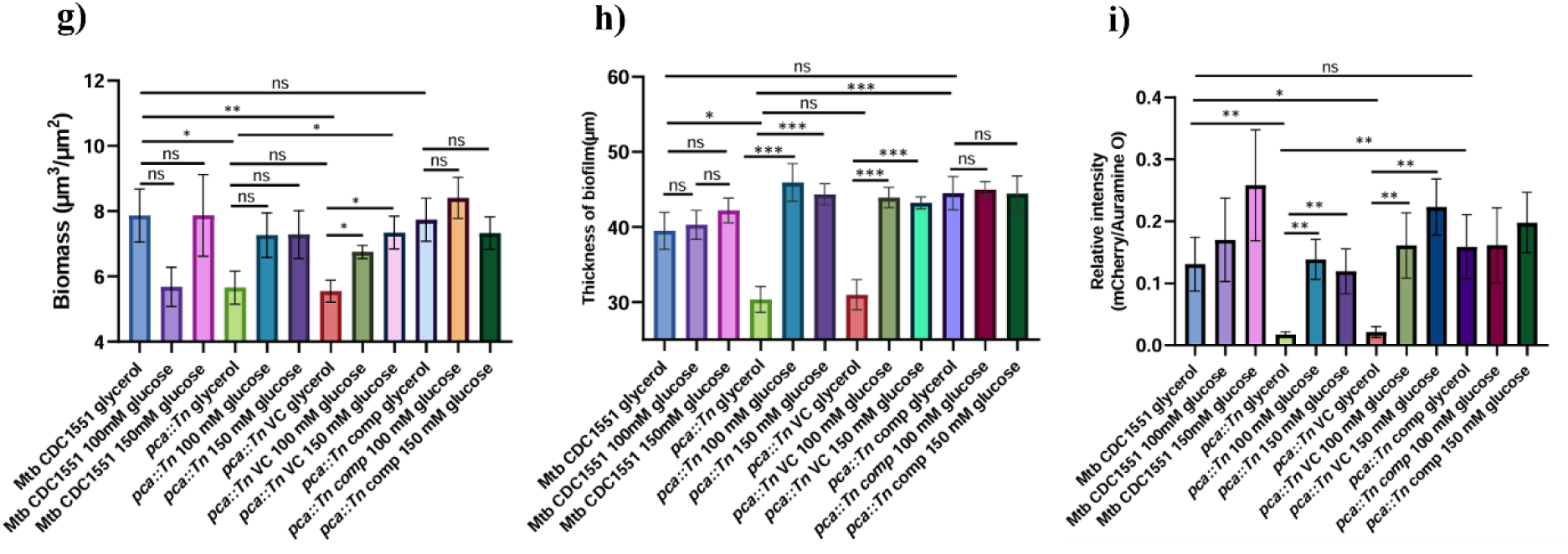
The addition of glucose regains the formation of thick, voluminous submerged biofilm structures in the *pca::Tn* strain when visualized by CLSM. Representative 3D images of submerged biofilm of all four strains i.e., Mtb CDC1551, *pca::Tn*, *pca::Tn VC*, and *pca::Tn comp* a) under glycerol-fed conditions. **b)** under 100 mM glucose-fed conditions. **c)** under 150 mM glucose-fed conditions. Representative side view images of submerged biofilm of all four strains **d)** under glycerol-fed conditions. **e)** under 100 mM glucose-fed conditions. **f)** under 150 mM glucose-fed conditions. **g)** Biomass (in µm^3^/µm^2^), **h)** thickness (in µm), and **i)** relative intensity analysis of the biofilm structures. Biomass and thickness analysis was performed through the COMSTAT plugin in ImageJ software, and relative intensity was calculated through IMARIS version 8. software. Data represent mean±SEM from three independent experiments. GraphPad Prism 8 was used to plot the column bar graphs. Statistical significance was determined using Student’s t-test, * indicates a p-value of <0.05, ** indicates a p-value of <0.005, *** indicates a p-value of <0.0005.

### Complementation of the *pca* gene or addition of glucose leads to restoration of the defect in colony phenotype in the *pca::Tn* strain

After observing the defect in biofilm formation in the *pca::Tn* mutant strain, we sought to analyze whether the defect could be restored in the 7H11OADC and 7H11OADC CR-CBB media plates. For this experiment, we took Mtb CDC1551, *pca::Tn*, *pca::Tn VC*, and *pca::Tn comp* strains and performed spot assay on 7H11OADC containing glucose as the sole carbon source and plate containing glycerol as a control **(Fig. 8a)**. We observed that the mutant strain regained the growth defect, and restoration of cording in colony morphology was visible upon the addition of 100 mM and 150 mM glucose **(Fig. 8b, c)**. In addition, the complemented strain grew at a rate similar to the wild type on control and glucose plates. Along with these, when the experiment was replicated onto 7H11OADC CR-CBB plates, we observed that *pca::Tn*, which is otherwise defective in color uptake, regains the phenotype upon glucose addition, and the complemented strain has the capability of color uptake in control as well as the glucose-supplemented plates behaving similar to wild type strain **(Fig. 8d-f)**. These results suggested that glucose plays a pivotal role in the biofilm formation and colony morphology of mycobacterial strains.

**Fig. 8:**
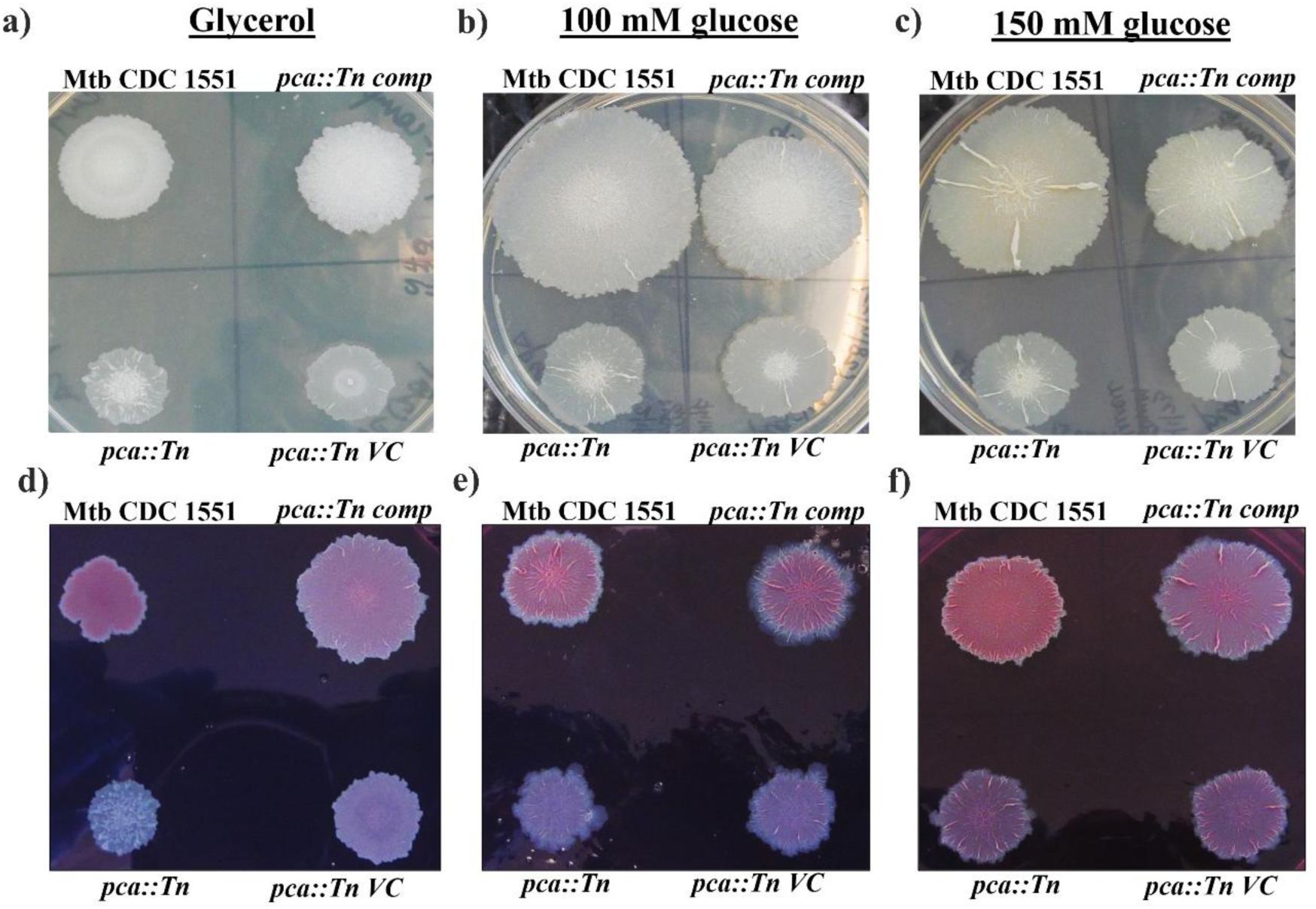
The addition of glucose or *pca* gene complementation restores the colony morphology phenotype in the mutant strain. Differences in colony morphology between Mtb CDC1551, *pca::Tn*, *pca::Tn VC*, and *pca::Tn comp* strains on 7H11OADC plates **a)** containing glycerol as a carbon source **b)** containing 100 mM glucose as a carbon source **c)** containing 150 mM glucose as a carbon source Differences in colony morphology between Mtb CDC1551, *pca::Tn*, *pca::Tn VC*, and *pca::Tn comp* strains on 7H11OADC CRCBB agar **d)** containing glycerol as a carbon source **e)** containing 100 mM glucose as a carbon source **f)** containing 150 mM glucose as a carbon source. Images are representative of three independent experiments.

### Complementation of the *pca* gene or addition of pyruvate leads to restoration of the defect in pellicle biofilm phenotype in the *pca::Tn* strain

In addition to the glucose, we also checked whether the phenomenon replicates upon pyruvate supplementation. Pyruvate at conc. 20 mM, 50 mM, and 70 mM were supplemented in the Sauton’s media, and glycerol was used as a positive control. Pyruvate was able to supplement the biofilm formation phenotype onto the *pca::Tn* mutant phenotype, and the addition of pyruvate led to increased biofilm formation with increasing concentrations **(Fig. 9a, b)**. However, the biofilm structure formed with the pyruvate supplementation was fragile and lacked significant cordings compared to the wild-type strain grown in glucose supplementation **(Fig. 5e, f)**. These data were quantitated using CV assay. The CV assay values were plotted, and the graph obtained showed a significant difference between the Mtb CDC1551 and *pca::Tn* mutant strain when grown upon glycerol as a carbon source. As the pyruvate is supplemented, the mutant strain showed biofilm formation comparable to the wild-type strain. However, the pyruvate supplementation is not as effective as the glucose supplementation as the concentrations could not match the complementation to the control/glycerol WT panel **(Fig. 9c)**.

**Fig. 9:**
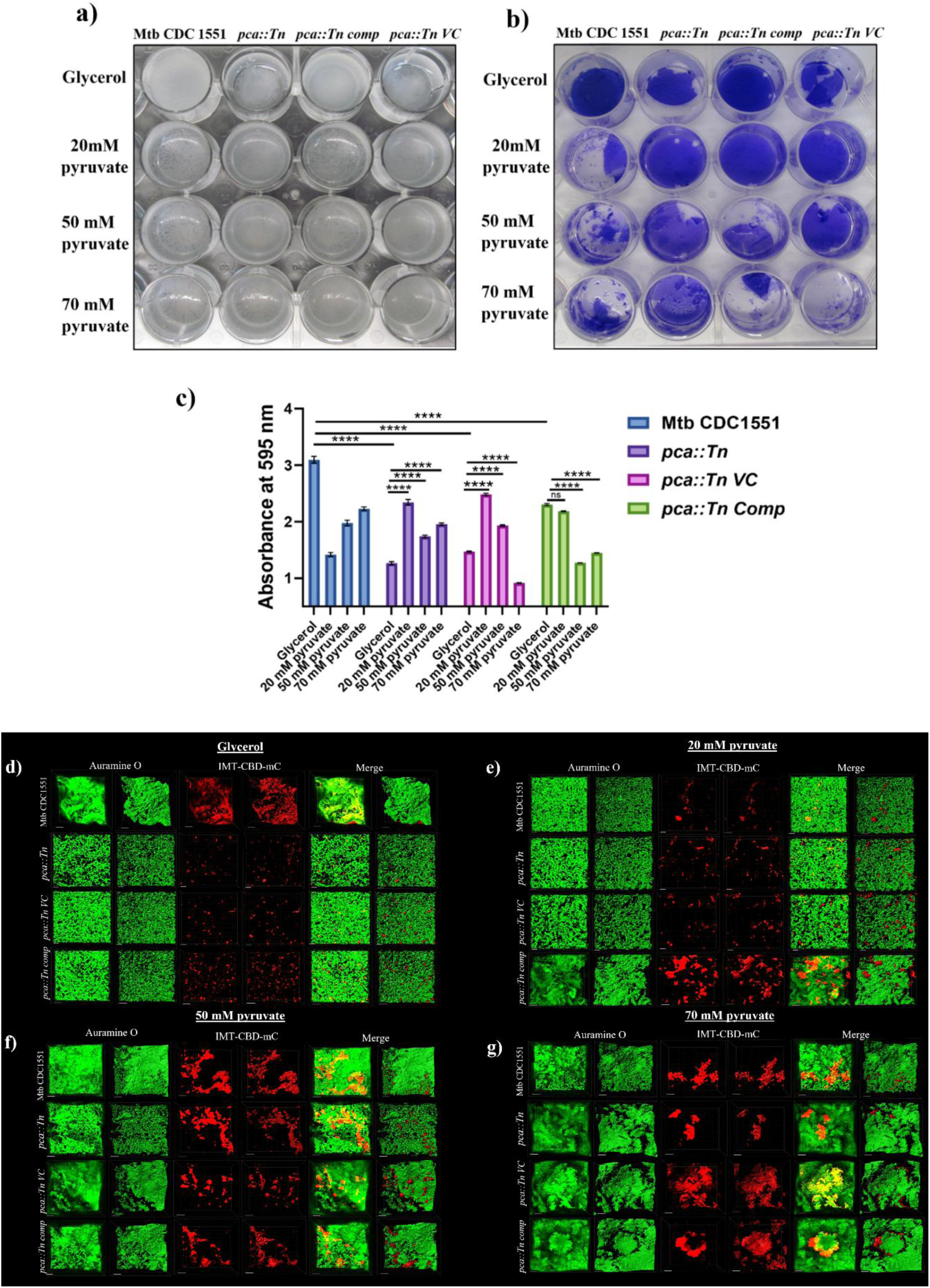
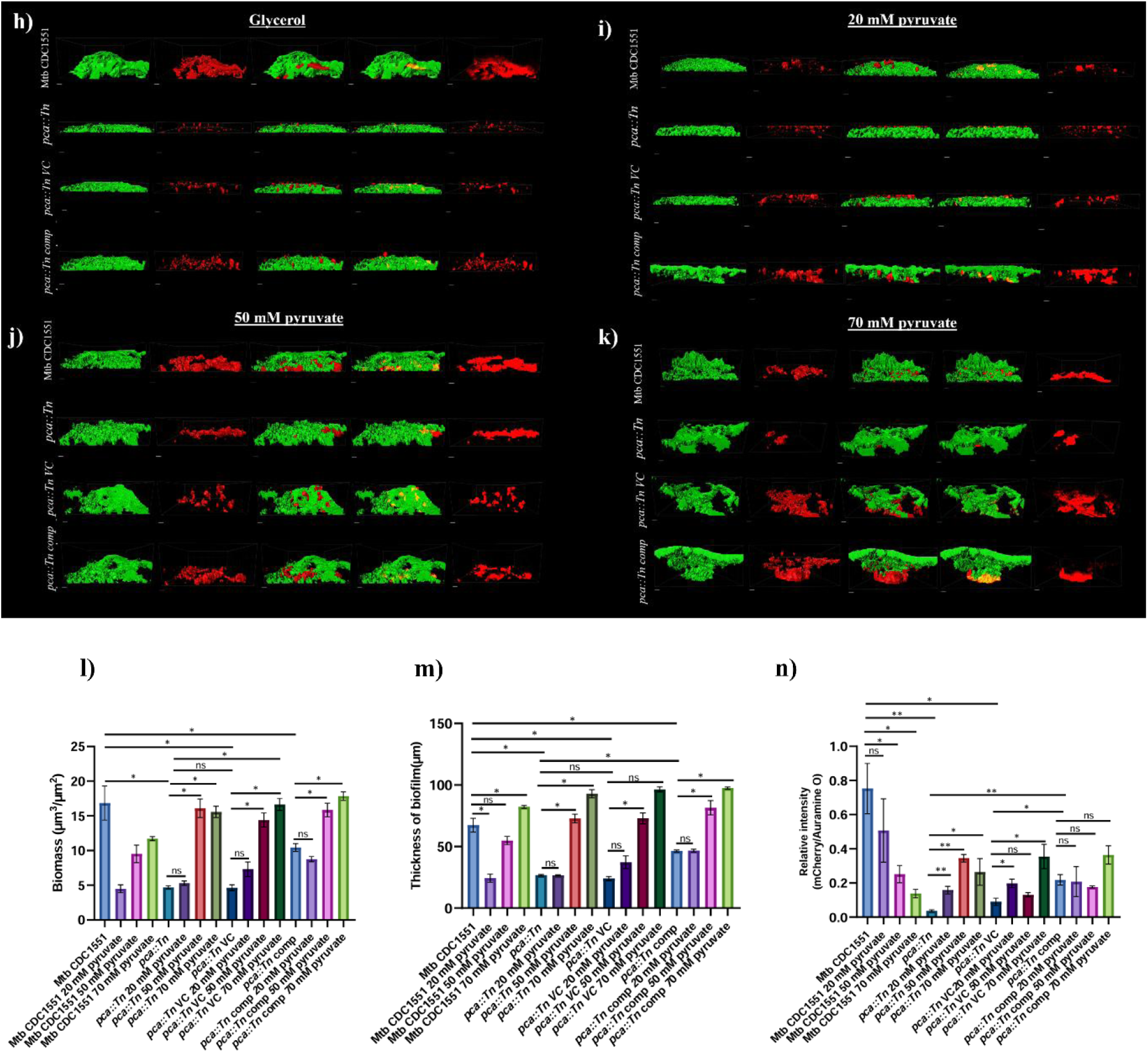
The addition of pyruvate regains the formation of thick, voluminous pellicle biofilm structures in the *pca::Tn* strain when visualized by CLSM. a) Representative image of pellicle biofilm formation in 24-well plates of Mtb CDC1551, *pca::Tn*, *pca::Tn VC*, and *pca::Tn comp* strains with pyruvate and glucose as carbon sources. **b)** Representative image of CV assay performed on pellicle biofilm setup. **c)** Graph showing CV assay of pellicle biofilm of Mtb CDC1551, *pca::Tn*, *pca::Tn VC*, and *pca::Tn comp* strains with pyruvate and glucose as carbon sources. Representative 3D images of pellicle biofilm of all four strains **d)** under glycerol-fed conditions, **e)** under 20 mM pyruvate-fed conditions, **f)** under 50 mM pyruvate-fed conditions, **g)** under 70 mM pyruvate-fed conditions. Representative side view images of pellicle biofilm of all four strains **h)** under glycerol-fed conditions, **i)** under 20 mM pyruvate-fed conditions, **j)** under 50 mM pyruvate-fed conditions, **k)** under 70 mM pyruvate-fed conditions. **l)** Biomass (in µm^3^/µm^2^), **m)** thickness (in µm), and **n)** relative intensity analysis of the biofilm structures. Biomass and thickness analysis was performed through the COMSTAT plugin in ImageJ software, and relative intensity was calculated through IMARIS version 8. software. Data represent mean±SEM from three independent experiments **(c, l, m, n)**. Statistical significance was determined using Two-way ANOVA, **** indicates a p-value of <0.0001 **(c)**. Statistical significance was determined using Student’s t-test, * indicates a p-value of <0.05, ** indicates a p-value of <0.005, *** indicates a p-value of <0.0005, **** indicates a p-value of <0.0001 **(l, m, n)**. GraphPad Prism 8 was used to plot the column bar graphs.

In addition to quantification via CV assay, we also imaged the pellicle biofilm formed by the above-mentioned strains via CLSM in the presence of glucose and pyruvate. We have stained the different mycobacterial strains using Auramine O dye and cellulose using an IMT-CBD- mC probe, which specifically binds to the cellulose through CBD. Here, we observed that in the case of glycerol as a given carbon source, the Mtb CDC1551 strain showed thicker and fully mature biofilms in contrast to the *pca::Tn* and *pca::Tn VC* strain where the biomass in the biofilm structure was lesser than in comparison with Mtb CDC1551 and *pca::Tn comp* strain and also consisted of few gaps/spaces in between the biofilm structures **(Fig. 9d)**. In the case of pyruvate supplementation, at 20 mM, we observed that all four strains formed thin biofilms with less biomass, with Mtb CDC1551 lacking the well-formed 3D biofilm **(Fig. 9e)**. Whereas at 50 mM and 70 mM conc. all the strains regained the phenomenon and formed thicker biofilms similar to the Mtb CDC1551 wild-type strain with no gaps/spaces in the structure. We also observed that IMT-CBD-mC staining was diminished in the mutant strain in the case of the control panel, whereas increased staining was visualized in the case of 50 mM and 70 mM pyruvate panels for all four strains **(Fig. 9f, g)**. Representative images of the side view of the biofilm panels are shown in (**Fig. 9h-k)**, which showcases the biofilm thickness of different strains. Biomass and thickness analysis were performed for the panels described above, and statistical difference was observed between Mtb CDC1551 and *pca::Tn*, and *pca::Tn VC* and between the *pca::Tn* glycerol and *pca::Tn* pyruvate panels highlighting the role of pyruvate in biofilm formation **(Fig. 9l, m)**. Relative intensity was calculated between mcherry/Auramine O fluorescence to showcase the increasing mcherry intensity in panels with glucose concentrations **(Fig. 9n)**. Although 20 mM pyruvate was incapable of inducing biofilm formation in all the strains, it can be concluded that 50 mM and 70 mM pyruvate conc. can be used for supplementation in biofilm formation. This suggested that pyruvate may also help in biofilm formation in addition to glycerol and glucose.

### Complementation of the *pca* gene or addition of pyruvate leads to restoration of the defect in colony phenotype in the *pca::Tn* strain

After observing the biofilm formation defect in the *pca::Tn* mutant strain, we sought to check whether the defect could be restored in the 7H11OADC and 7H11OADC CR-CBB media plates. For this experiment, we took Mtb CDC1551, *pca::Tn*, *pca::Tn VC*, and *pca::Tn comp* strains, and performed spot assay on 7H11OADC containing pyruvate as the sole carbon source and plate containing glycerol as a control **(Fig. 10a)**. We observed that the mutant strain was able to restore the growth, cording and colony morphology defects upon addition of 20 mM, 50 mM, and 70 mM pyruvate **(Fig. 10b-d)**. In addition, the complemented strain was able to grow well, similar to the wild type on control as well as on pyruvate plates. Along with these, when the experiment was replicated onto 7H11OADC CR-CBB plates, where we observed that *pca::Tn*, which is otherwise defective in color uptake, regains the phenotype upon pyruvate addition, and the complemented strain has the capability of color uptake in control as well as the pyruvate supplemented plates behaving similar to wild type strain **(Fig. 10e-h)**. Although the complementation was not comparable to the glucose supplementation, as seen in **Fig. 8b, c, e,** and **f**. These results suggested that in addition to biofilm formation, pyruvate plays a key role in the colony morphology of mycobacterial TB strains.

**Fig. 10:**
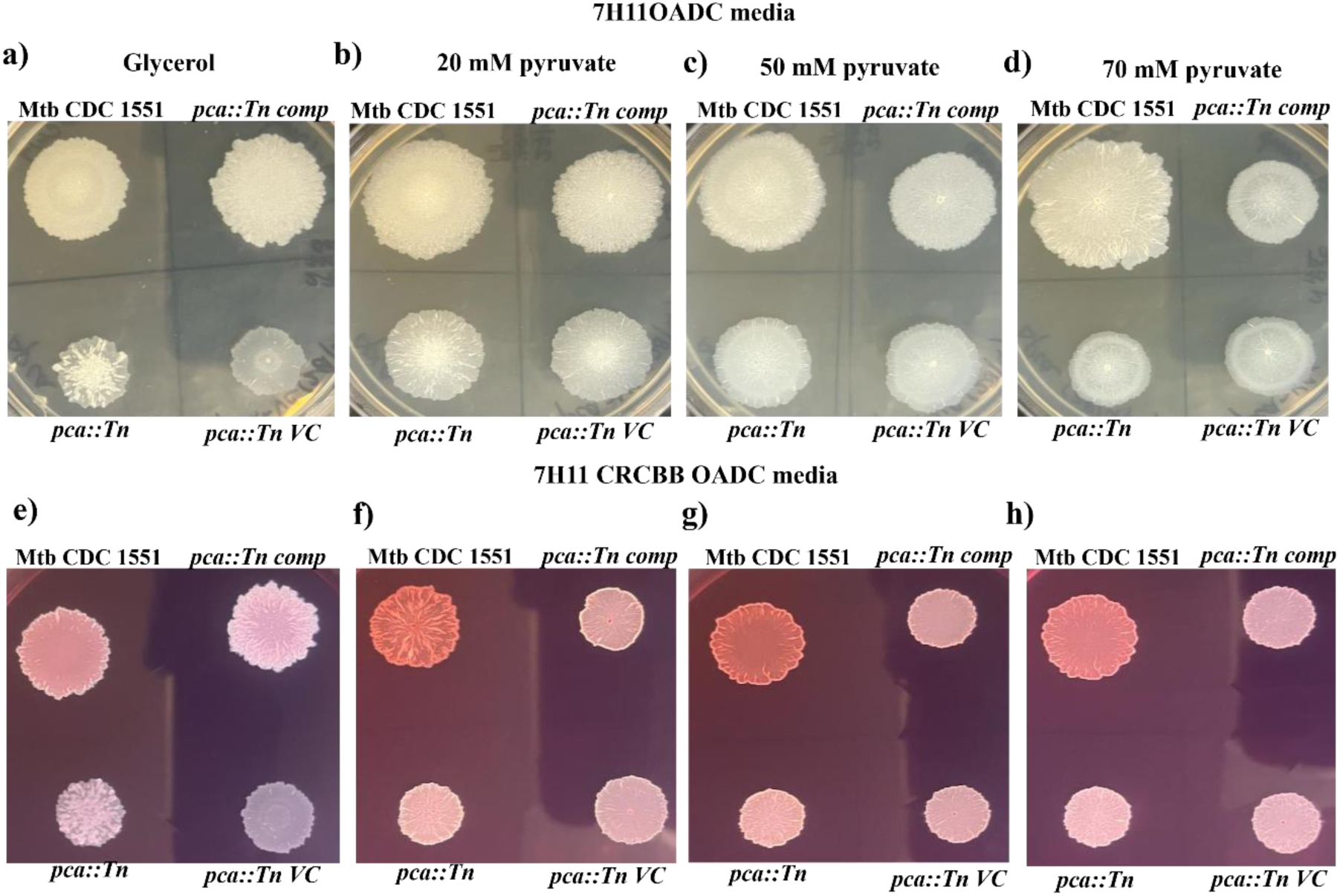
The addition of pyruvate or *pca* gene complementation restores the colony morphology phenotype in the *pca::Tn* mutant strain. Differences in colony morphology between Mtb CDC1551, *pca::Tn*, *pca::Tn VC*, and *pca::Tn comp* strains on 7H11 OADC plates **a)** containing glycerol as a carbon source **b)** containing 20 mM pyruvate as a carbon source **c)** containing 50 mM pyruvate as a carbon source **d)** containing 70 mM pyruvate as a carbon source. Differences in colony morphology between Mtb CDC1551, *pca::Tn*, *pca::Tn VC*, and *pca::Tn comp* strains on 7H11 OADC CR-CBB agar **e)** containing glycerol as a carbon source **f)** containing 20 mM pyruvate as a carbon source **g)** containing 50 mM pyruvate as a carbon source **h)** containing 70 mM pyruvate as a carbon source. Images are representative of three independent experiments.

## Discussion

The role of *pca* has been previously characterized in the intracellular survival of Mtb and the anaplerosis pathway. However, the role of gluconeogenesis and its key enzymes in biofilm formation has not been explored. In this study, we first checked the impact of different carbon sources on mycobacterial biofilm formation. Herein, we found that apart from glycerol (classical carbon source), glucose and pyruvate can also contribute to mycobacterial biofilm formation. Glucose replenished the biofilm phenomenon, similar to glycerol, and showed better complementation at higher concentrations. Pyruvate restored the phenomenon to a certain extent but not to the level of glucose/glycerol. We also utilized the *pca::Tn* mutant and checked its role in biofilm formation and growth under various carbon sources. Under normal growth conditions, it was observed that *pca::Tn* showed a delayed growth pattern in comparison to the WT strain and was found to be defective in pellicle biofilm formation. Upon its supplementation with glucose or its complementation with the gene sequence, *pca::Tn* restored the biofilm formation phenomenon, depicting its relevance in pellicle biofilm formation. In addition to the classical pellicle biofilm formation, we have also checked the role of *pca* in TRS-induced submerged biofilm formation. This biofilm model was discovered by Trivedi *et al.,* and has shown that biofilm EPS is rich in polysaccharides such as cellulose and other components, i.e., proteins, extracellular DNA, and lipids^15^. *pca::Tn* mutant also showed a defective submerged biofilm formation with gaps/spaces present in between the biofilm structure. This phenotype was complemented upon the superficial addition of different glucose concentrations in the medium or the *pca::Tn comp* (*pca* gene complemented) strain. We also visualized the pellicle and submerged biofilm via CLSM imaging and checked for the presence of cellulose in the biofilms using an IMT-CBD-mC probe designed by Mavi *et al*^13,14^. The presence of cellulose was observed via IMT-CBD-mC staining in Mtb CDC1551 and *pca::Tn comp* strains and enhanced upon increasing conc. of glucose and pyruvate. These results suggested that *pca* plays an important role in biofilm formation and modulates the polysaccharide content in pellicle and submerged biofilm models. Previous reports in microbes such as *P. aeruginosa* and *S. aureus* have shown the importance of the *pca* gene in biofilm formation. In *P. aeruginosa*, in addition to being involved in several metabolic pathways, this gene is shown to be important for pathogenicity and biofilm formation during *in vitro* growth^21^. In *S. aureus*, the absence of *pyc* (pyruvate carboxylase) gene led to decreased biofilm formation and was restored upon genetic complementation. It has also been shown to be involved in pathogenesis and required for intracellular survival^22^.

Through these results, it can be concluded that *pca* is important for biofilm formation. We hypothesize that as the *pca* feeds the carbon flux, *i.e.*, oxaloacetate, into the gluconeogenesis pathway, which then is required to generate glucose molecules **(Fig. 11)**. These glucose molecules are then utilized to generate energy or as a substrate for EPS production under biofilm-forming conditions. Recent research on *S. cerevisiae*, *S. aureus*, *S. enterica* serovar *Typhimurium*, *C. albicans*, *V. cholerae,* and others have demonstrated that gluconeogenesis intermediates are necessary for EPS synthesis and biofilm development^8–12^. Glucose molecules can then be used as EPS building blocks for generating a variety of polysaccharides. Since the biofilm models represented in this study show the presence of cellulose, confirmed via the IMT-CBD-mC probe staining, it can be suggested that the generation of cellulose is achieved through gluconeogenesis under biofilm development conditions. These observations suggest a critical role for the gluconeogenesis pathway in mycobacterial biofilm development, as the non-functionality of this pathway leads to defective biofilm formation. The model depicts the role of gluconeogenesis in mycobacterial biofilm development, illustrated in **Fig. 12**.

**Fig. 11:**
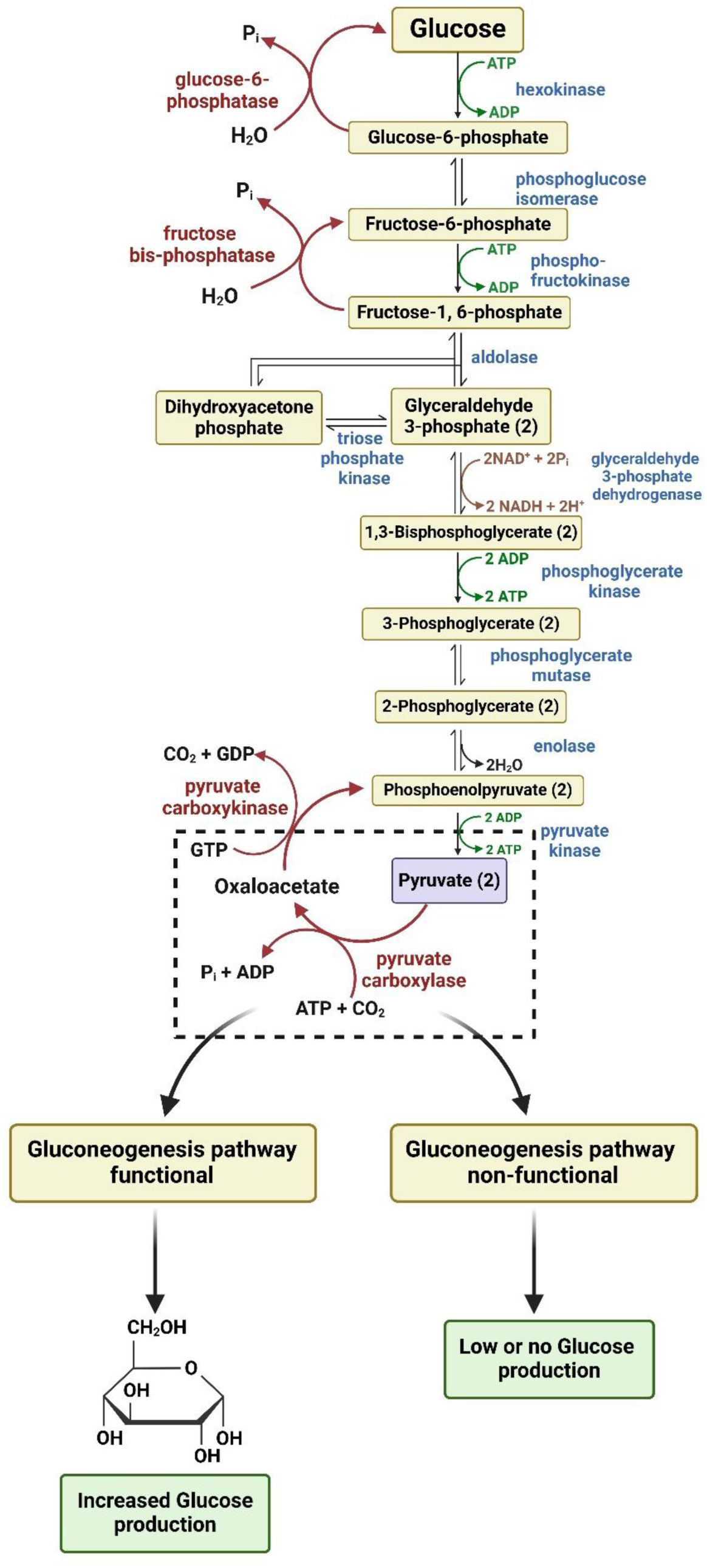
Pyruvate carboxylase facilitates the carbon flux into the gluconeogenesis pathway. Upon functional pyruvate carboxylase in Mtb, the carbon flux is fed into the gluconeogenesis pathway via non-carbohydrate sources such as oxaloacetate. The functional gluconeogenesis pathway ensures sufficient production of glucose molecules. Whereas, when the bacteria are deprived of pyruvate carboxylase, the gluconeogenesis pathway becomes non-functional, and low or no glucose production is observed, suggesting pyruvate carboxylase is essential for glucose levels in bacterial cells (Created in BioRender. Kumar, A. (2025) https://BioRender.com/c70b017).

**Fig. 12:**
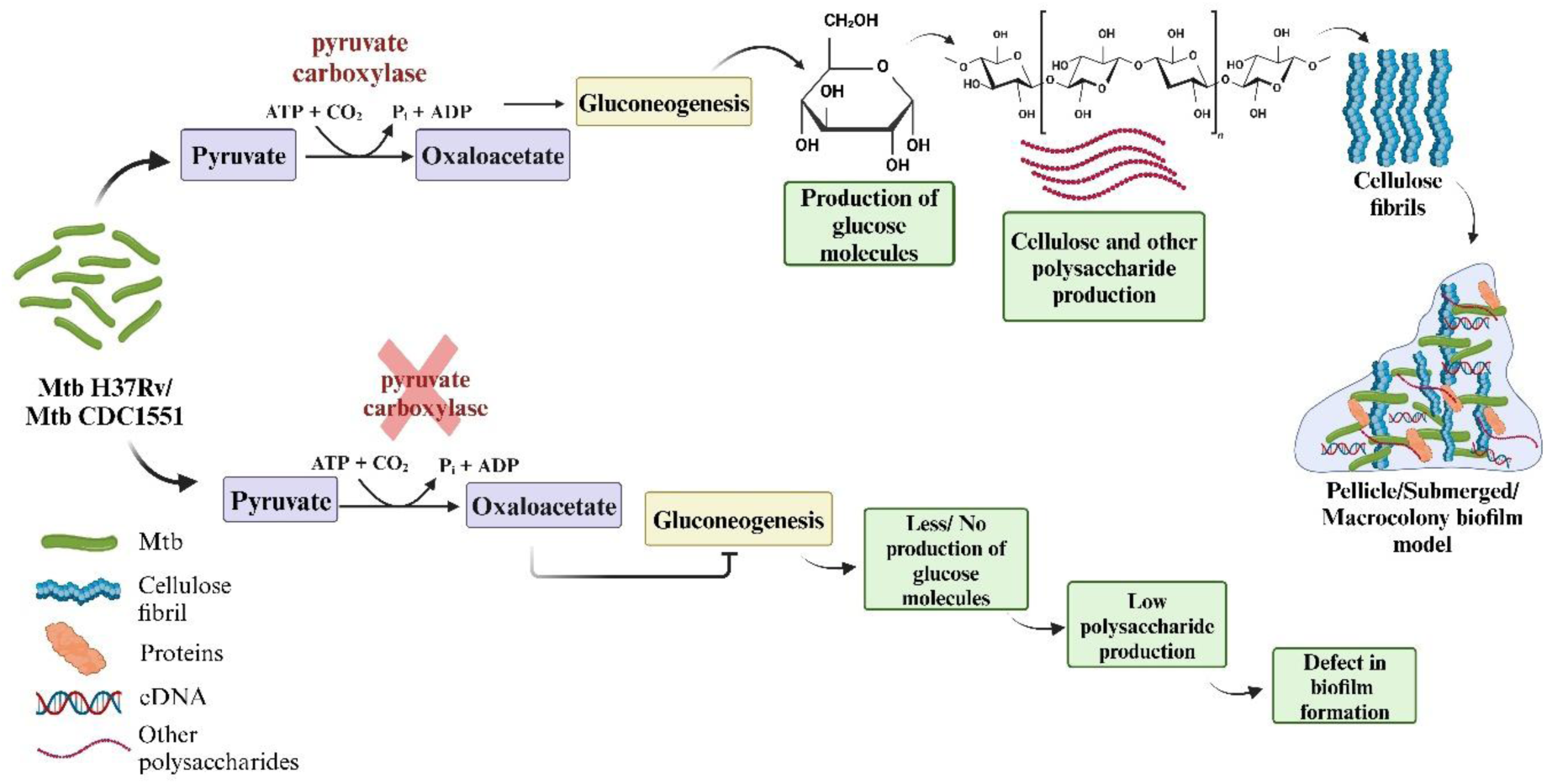
Gluconeogenesis plays a critical role in the formation of mycobacterial TB biofilm. Mtb H37Rv or Mtb CDC1551 strains, under normal conditions, are able to feed the carbon flux through the gluconeogenesis pathway, thereby generating glucose molecules. The generation of glucose molecules is required to produce cellulose and other polysaccharides whose primary subunit is glucose. These polysaccharides are required to integrate into the pellicle, submerged, and macrocolony biofilm models. The integration of cellulose contributes to the structural backbone of these biofilm models. Upon obstruction of the gluconeogenesis pathway, the generation of glucose molecules is restricted, and thereby, less/no production of glucose molecules occurs. This results in low polysaccharide production, thereby, defective pellicle, submerged, and macrocolony biofilm models. This hypothesis ensures the essentiality of the gluconeogenesis pathway in mycobacterial biofilm formation (Created in BioRender. Kumar, A. (2025) https://BioRender.com/d39e222).

## Materials and Methods

### Bacterial medium and culturing conditions

Mtb H37Rv (ATCC 27294), Mtb CDC1551 (Bei Resources, NR-13649), Mtb CDC1551 Tn Mutant 2242 (MT3045, *rv2967c*) (Bei Resources, NR-15722) were grown in complete 7H9 broth [7H9 (Becton Dickinson Difco, 271310) supplemented with 0.2% glycerol (Rankem, G0040), 0.1% Tween 80 (MP, 103170) and 10% (OADC)]. The vector control strain pstHIT vector in Mtb CDC1551 Tn Mutant 2242, i.e., *pca*::*Tn VC,* and complementing strain containing pstHIT+*pca* gene i.e., *pca::Tn comp* were grown in Middlebrook 7H9 broth or solid Middlebrook 7H11 OADC agar along with additional hygromycin (50 μg/ml). The cultures were incubated at 37°C, 90 rpm in 30 mL or 60 mL square polyethylene terephthalate glycol media bottles until the desired OD_600_ was reached. 7H11 agar (BD Difco, 212203), along with 0.5% glycerol (Rankem, G0040) and 10% OADC, was used as solid media.

For growth curve experiments, exponentially growing cultures of Mtb were washed and inoculated in 7H9 medium with all the supplements to achieve an initial OD_600_ of 0.1. Their growth rate was recorded for 22 days until they reached the stationary phase of growth. All the experiments utilizing Mtb were carried out inside the institute’s BSL-3 facility in accordance with the standard operating procedures.

### Construction of *pca* complemented strain

Cloning was carried out using KOD polymerase (Toyobo, KOD-201) and restriction digestion with Fast Digest range of enzymes from Thermo Scientific (EcoRV; FD0304 and HindIII; FD0504). The gene *rv2967c* was amplified using primers enlisted in **Table 1**. and was cloned in a mycobacterial integration vector, pstHIT, with a hygromycin gene cassette used as a selection marker. N-terminal his-tag was already present in the vector map. Finally, it was transformed in Mtb CDC1551 Tn Mutant 2242 (MT3045, *rv2967c*) to generate the complementing strain, and the alone vector without the gene sequence was also transformed to create a vector control strain. For the qRT-PCR experiment, RNA was isolated using a Direct-Zol RNA isolation kit (ZYMO). The primers used for RT-qPCR to detect mRNA levels are enlisted in **Table 2**. cDNA synthesis and PCR reactions were carried out using total RNA extracted from each bacterial culture and Superscript III platinum-SYBR green one-step qRT-PCR kit (Invitrogen) with appropriate primer pairs (2 µM) using an ABI real-time PCR detection system. Endogenously expressed Mtb *rpoB* was used as an internal control. The ΔΔCT method calculated the fold difference in gene expression^23^.

**Table 1.**
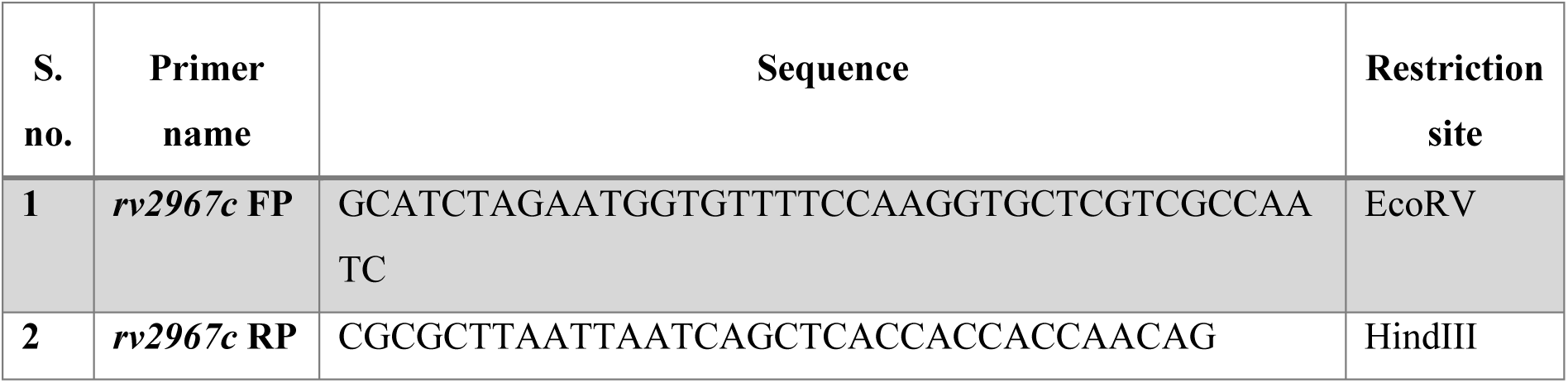
List of primers used for creating a complementing strain of the *rv2967c* (*pca)* gene.

**Table 2.**
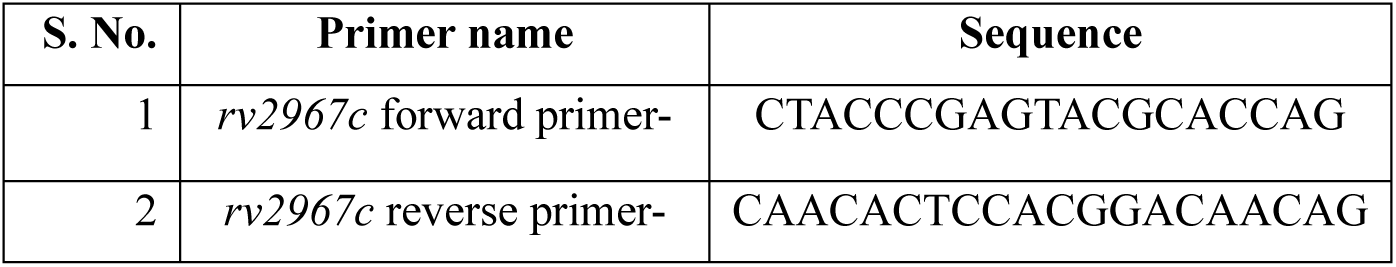
List of oligonucleotide primer sequences used in RT-qPCR to detect mRNA levels in the mutant and complemented strains.

### Pellicle biofilm assays and colony morphology

For pellicle biofilm formation, saturated cultures of Mtb CDC WT, *pca::Tn*, *pca:: Tn VC,* and *pca::Tn comp* strains were inoculated (10% inoculum) in 2 ml of Sauton’s media in 24 well plates. Pellicle biofilms were formed in 8-well-chambered glass slides for CLSM. A 10% inoculum was added in 500 µl of Sauton’s media in 8-well chambered glass slides and incubated for approximately 3-4 weeks at 37°C in standing conditions. To form submerged biofilms, logarithmic-phase cultures of Mtb CDC WT, *pca::Tn*, *pca::Tn VC*, and *pca::Tn comp* strains (grown in 7H9 media+5% OADC) were induced with 6 mM DTT for 29 hr at 37°C in 24 well plates. Submerged biofilms were also formed in 8-well-chambered glass slides for CLSM. For inoculation in 8-well chambered glass slides, cultures of OD_600_ 0.8–1 were taken, 500 µL was transferred to each well and was induced with 6 mM DTT.

### CV assay of biofilms

The CV assay of Mtb biofilms was performed as described earlier^15^. Briefly, the CV assay was performed in 24-well plates. After forming the Mtb pellicle or submerged biofilm (in 24 well plates), the media was removed and washed with 1X phosphate buffer saline (PBS), and 2 ml of 1% CV (sigma) was gently added to the biofilm. It was incubated for 20 min at 37°C. The stain was removed, and the biofilm was gently washed twice with 1XPBS. The bound CV was then extracted by a 10-minute incubation at 37°C with 2 ml of 95% ethanol. The absorbance of extracted CV was measured at 595 nm on a spectrophotometer. The CV assay was performed in three independent replicates for each dataset.

### Preparation of plates containing CR and CBB dye, containing glucose and pyruvate

For the construction of 7H11OADC, media plates containing glycerol were prepared as described previously^13^. Glycerol was eliminated from the 7H11 OADC media plates and was instead replenished with glucose and pyruvate. For glucose 100 mM and 150 mM conc. of dextrose (Merck) were added as per the required media volume. Similarly, for pyruvate, 20 mM, 50 mM, and 70 mM conc. of sodium pyruvate (Sigma) was added as per the required media volume. For 7H11OADC CR-CBB media plates, media was supplemented with CR (40 μg/ml), and CBB (20 μg/ml) along with the respective carbon sources.

For spot assay, 2 μl of each culture logarithmic cultures with similar OD_600_ were spotted onto solid Middlebrook 7H11 OADC and 7H11 OADC CRCBB agar containing either glycerol, glucose or pyruvate as a carbon source. The plates were incubated at 37°C for approximately four weeks.

### Confocal laser scanning microscopy of pellicle and submerged biofilms

Mature pellicle and submerged Mtb biofilms were produced on chambered glass slides for CLSM imaging. After the maturation of biofilms, the sample was fixed with 4% paraformaldehyde for 1 hour and washed thrice with 1XPBS. After washing, it was stained with fluorescent probes such as Auramine O (for Mtb staining), and IMT-CBD-mC (for cellulose) (10 µM). Biofilms were stained with Auramine O for 15 min, and IMT-CBD-mC was used as a stain at 10 µM concentration in TRIS buffer (50 mM, pH 7.5) with 1 M NaCl for 1-1.5 hrs. After staining, samples were washed thrice with PBS. Stained biofilms were viewed using a Nikon A1R confocal microscope.

## Statistical analysis

Results are expressed as mean ± SEM from at least three biological replicates. Statistical significance was determined using GraphPad Prism 8 employing students’ t-test (unpaired, non-parametric Gaussian distribution, Mann-Whitney test) and Two-way ANOVA. * depicts *p* value <0.05, ** depicts *p* value <0.01, *** depicts *p* value <0.001, and **** *p* value <0.0001.

## Acknowledgments

AK is supported through Grant No. IA/S/20/2/505220 by DBT/Wellcome Trust India Alliance (India Alliance). SS and SB are supported by a fellowship under the India Alliance project. Illustrations of this manuscript were created with BioRender.com.

## Author Contributions

SS and SB designed and performed the experiments. SS performed the analysis of the experiments. SS wrote the original draft, and AK and SS reviewed the manuscript. AK conceived and coordinated the study.

## Data availability statement

All the data pertaining to the results reported in this study are part of the main text.

## Abbreviations

CBD: Cellulose binding domain
CLSM: Confocal laser scanning microscopy
CR: Congo Red
CBB: Coomassie Brilliant Blue dyes
CV: Crystal violet
DTT: Dithiothreitol
EPS: Extracellular polymeric substances
Mtb: Mycobacterium tuberculosis
OADC: Oleic acid-albumin-dextrose-sodium chloride
PBS: Phosphate buffer saline
*pckA*: Pyruvate carboxykinase
*pca*: Pyruvate carboxylase
TRS: Thiol reductive stress
Tn: Transposon

